# Capturing limbal epithelial stem cell population dynamics, signature, and their niche

**DOI:** 10.1101/2020.06.30.179754

**Authors:** Anna Altshuler, Aya Amitai-Lange, Noam Tarazi, Sunanda Dey, Lior Strinkovsky, Swarnabh Bhattacharya, Shira Hadad-Porat, Waseem Nasser, Jusuf Imeri, Gil Ben-David, Beatrice Tiosano, Eran Berkowitz, Nathan Karin, Yonatan Savir, Ruby Shalom-Feuerstein

## Abstract

Stem cells (SCs) are traditionally viewed as rare, slow-cycling cells that follow deterministic rules dictating their self-renewal or differentiation. It was several decades ago, when limbal epithelial SCs (LSCs) that regenerate the corneal epithelium were one of the first sporadic, quiescent SCs ever discovered. However, LSC dynamics, heterogeneity and genetic signature are largely unknown. Moreover, recent accumulating evidence strongly suggested that epithelial SCs are actually abundant, frequently dividing cells that display stochastic behavior.

In this work, we performed an in-depth analysis of the murine limbal epithelium by single-cell RNA sequencing and quantitative lineage tracing. The generated data provided an atlas of cell states of the corneal epithelial lineage, and particularly, revealed the co-existence of two novel LSC populations that reside in separate and well-defined sub-compartments. In the “outer” limbus, we identified a primitive widespread population of quiescent LSCs (qLSCs) that uniformly express Krt15/Gpha2/Ifitm3/Cd63 proteins, while the “inner” limbus host prevalent active LSCs (aLSCs) co-expressing Krt15-GFP/Atf3/Mt1-2/Socs3. Analysis of LSC population dynamics suggests that while qLSCs and aLSCs possess different proliferation rates, they both follow similar stochastic rules that dictate their self-renewal and differentiation. Finally, T cells were distributed in close proximity to qLSCs. Indeed, their absence or inhibition resulted in the loss of quiescence and delayed wound healing. Taken together, we propose that divergent regenerative strategies are tailored to properly support tissue-specific physiological constraints. The present study suggests that in the case of the cornea, quiescent epithelial SCs are abundant, follow stochastic rules and neutral drift dynamics.

## Introduction

Stem cells (SCs) differ from any other cells by their substantial ability to retain in an undifferentiated state, self-duplicate or enter differentiation program, depending on tissue demand^1 2^. SCs have been successfully applied in clinical trials to reconstitute the bone marrow, treat skin burn, and restore corneal blindness^3^. However, various fundamental features of SCs are still under debate or remain unknown. For example, SC prevalence in the niche, SC heterogeneity and self-renewal mechanisms are not well understood, and consequently, they remain enigmatic cells^4 5^. These key challenges hamper the progression towards advanced application of SCs for regenerative medicine.

Seminal studies supported a deterministic and hierarchical model in which SCs were viewed as slow-cycling cells that are located in a specialized microenvironment known as the niche^6 7 8^. Such quiescent SCs (qSCs) are commonly believed to be rare cells that are surrounded by their progeny, of abundant, fast cycling but short-lived progenitor cells^9^. Key evidence for the quiescence and scarcity of SCs came from indirect observations *in vivo* and *in vitro*. Sporadic slow-cycling cells were identified in different tissues as nucleotide-label retaining cells in pulse-chase experiments performed in various tissues including the cornea^10^, bone marrow^11^, brain^12^, skin^13^, gut^14^ and skeletal muscle^15^. Clonal survival/growth dynamics *in vivo* (i.e. by genetic lineage tracing of single cells at the SC compartment) or ex vivo (i.e. by colony formation assay), typically result in the detection of a low number of clones that survive for long-periods of time. This evidence shaped the traditional deterministic model that considered qSCs as very potent cells that are also very scarce. The benefits of quiescence may be paramount and include reduced biochemical damage associated with the production of toxic agents or biosynthesis of macromolecules, and minimalized accumulation of disease-causing mutations^16 17^.

Despite much effort in the field, markers that reliably identify genuine and scarce qSCs were usually not found. Proposed markers, typically labeled a contiguous widespread cell population in the SC compartment^5^. Nevertheless, the co-existence of qSCs and actively dividing SCs (aSCs) in specific tissues including the bone marrow, brain, hair follicle and muscle has been shown^18 19 20^. These tissues differ from traditional epithelial SC models by their significantly slower turnover (or resting phase)^21^. In recent years, however, frequently dividing SCs were identified in epithelial tissues including the gut^22^, epidermis^23^ and esophagus^24^. These studies proposed an alternative, virtually opposing, stochastic model that attributes SCs with entirely contrary properties including short cell cycle, abundance in the niche and unpredictable survival rate^18 1 25 26^. One of the common models that is used to capture this phenomena is that of asymmetric self-renewal with neural drift dynamics^23 27^. These findings challenged the conventional paradigm that considered quiescence as an integral feature of SCs.

It is likely that the appropriate regenerative strategy is tailored for each tissue depending on the physiological necessities and challenges (e.g. irradiation, toxicity, cellular turn over or lineage diversity). In this sense, the corneal epithelium is an interesting study case, whereas SCs and their progeny must support tissue transparency and protection from sun irradiation, toxicants, physical injury and invading microorganisms^28^. Indeed, most ocular surface epithelial cancers are restricted to the corneal SC compartment of the limbus. However, the rareness of limbal tumors suggests the existence of specialized mechanisms that may provide protection against neoplastic cell transformation.

The cornea is an excellent model for SC research hallmarked by segregation of SCs, short-lived progenitors and differentiated cells into distinct anatomical compartments. The clarity and high accessibility of the cornea, allows multi-color (“Confetti”) fluorescent lineage tracing^29 30 31^, assessing different SC hypotheses^32^, as well as SC or progenitor cell depletion using vital microscopy^33 34^. Corneal epithelial SCs reside in the limbus, a ring-shaped region at the corneal-conjunctival boundary. The limbal microenvironment (niche) markedly differs from that of the cornea. It contains blood and lymph vasculature whereas the cornea is avascular, it is hallmarked by a unique extracellular matrix with soft biochemical properties while it hosts specialized niche cells^35 36 37^.

Rare label retaining cells were confined to the limbus^10 38^. In line, colony formation tests revealed that only a low percentage of limbal cells could form large clones (“holoclones”) *in vitro*, a feature that is commonly attributed to bona fide limbal SCs (LSCs)^39 40^. Based on these observations, it is widely believed that true LSCs are scarce limbal epithelial cells kept most of the time in quiescence state and are surrounded by their early progeny which are abundant, fast dividing but short-lived progenitor cells.

Tremendous efforts have been made by many research groups to discover markers for the identification of LSCs. Keratin 15 (Krt15) ^33 41^, C/EBPdelta and BMI1^42^, ABCB5^43^, ABCG2^44^ and P63^45^ represent a partial list of genes proposed to identify LSCs. Recently, we have shown that the Krt15-GFP transgene (green fluorescent protein coding gene under the promoter of Krt15) labeled a discrete population of murine LSCs. Krt15-GFP^+^ basal limbal epithelial cells were located at close proximity to the site of corneal regeneration origin, as evident by lineage tracing of Krt14^+^ cells^33^.

However, neither Krt15-GFP, nor any of the proposed markers could faithfully mark a hypothetical rare, slow-cycling SC population in the limbus, suggesting that the current model may be inaccurate. Consequently, the prevalence of LSCs, their genetic signature, and regulation by niche cells remain poorly defined. Here we combined single-cell transcriptomics and quantitative lineage tracing to capture LSC populations and their signature. This work provides a useful atlas of the entire murine corneal epithelial lineage, including the signature of a slow cycling quiescent LSC (qLSC) state, and finally, unravels an apparently tight interaction and regulation of LSCs by T cells.

## Results

Careful inspection of the limbus of Krt15-GFP transgene bearing mice showed that Krt15-GFP^+^ cells are confined to an “inner” limbal zone, suggesting that the “outer” limbal region (the space between Krt15-GFP^+^ cells and the Krt8^+^ conjunctiva) contains a different LSC population (Fig. 1a, S1a). To clarify the molecular nature and heterogeneity of LSC populations, we performed a single-cell transcriptomic analysis. In order to facilitate the detection of rare cell populations, reduce variability and gain robust statistical data, we preferentially isolated epithelial cells from the limbus of 5 individual adult mice (2.5 months old, n=10 eyes). The limbus together with marginal conjunctiva and corneal periphery were carefully dissected (Fig. 1a), a protocol for isolating mainly epithelial cells was applied (see Methods), and suspended cells were subjected to 10x Chromium Single-Cell RNA sequencing (scRNA-seq). Initial raw data analysis and quality controls were performed with Cell Ranger software and a large number of 4,787 high quality cells passed rigorous quality tests exhibiting significant mean reads (98,763 per cell) as well as a substantial median number of 2,479 detected genes per cell.

**Figure 1:**
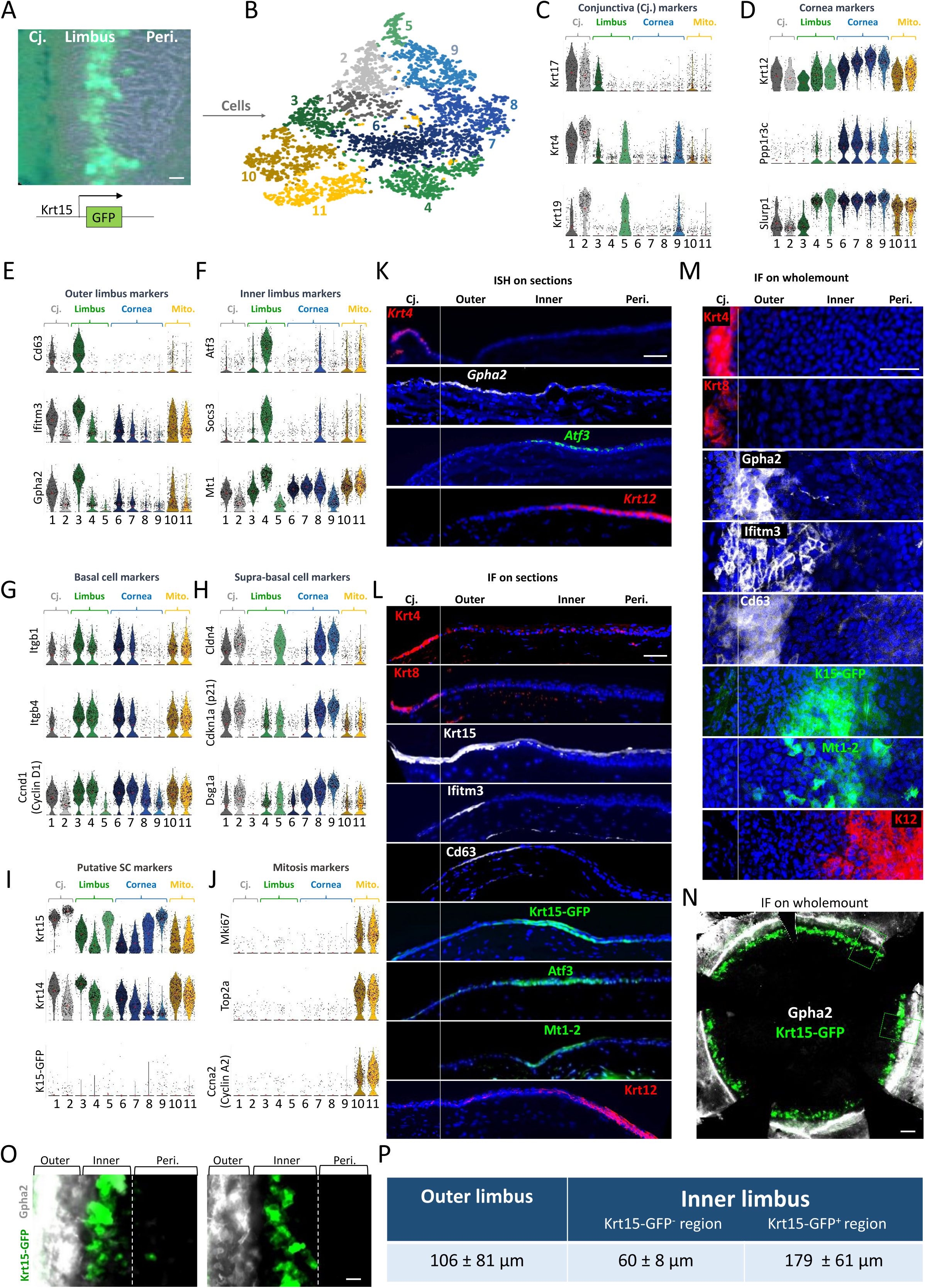
scRNA-seq reveals corneal epithelial cell states including 2 distinct limbal epithelial cell populations. (A-B) The limbus (with marginal conjunctiva and cornea) of 10 corneas of 2.5-month old Krt15-GFP transgenic mice was dissected, pooled and epithelial cells were sublected to scRNA-seq analysis. (B) t-SNE plot revealed 11 distinct clusters of cells. (C-J) Violin plots showing the expression of differentially expressed markers of specific clusters of cells that belong to the conjunctiva (Cj.), limbus, cornea or cells of mixed lineages that were captured during mitosis (Mito). (K-O) In situ hybridization (ISH) (K) or immunofluorescent staining (IF) (L) or whole mount staining (M-O) of the indicated markers performed on cornea tissue sections of 2-month old mice. The conjunctiva (Cj.) limbal compartments (outer or inner) and corneal periphery (Peri.) are indicated. (P) Size analysis (mean ± standard deviation) of the different limbal sub-compartments. Scale bars was 50 μm (A-M, O-P) or 300 μm (N). Nuclei were counter stained by DAPI. Abbreviations: Cj., Conjunctiva; Peri., Periphery.

### In silico analysis revealed discrete cell states in the corneal epithelial lineage

Cell filtering and unbiased clustering were performed with R package Seurat^46^ revealing 11 cell populations displaying a markedly distinct signature of gene expression (Fig. 1b, S1b). Analysis of lineage-specific markers suggested that >99% of the cells are ocular epithelial cells (see Methods). Next, we performed in silico analysis of putative markers for each epithelial tissue (i.e. conjunctiva, limbus, and cornea) and cell layer (i.e. basal or supra basal layer) (Fig. 1c-j, S1c-d).

#### Conjunctival cells

Clusters 1-2 were identified as conjunctival cells based on the positive expression of putative conjunctival cytokeratins (Krt17, Krt4, Krt19 (Fig. 1c) and Krt6a, Krt13, Krt8 (Fig. S1c)^47^), and the lack or comparatively low expression of corneal markers Krt12 and Slurp1^48 49^ and of the protein phosphatase 1 (Ppp1r3c) (Fig. 1d). Further analysis of putative markers of basal (Itgb1, Itgb4, Ccnd1 (Cyclin D1)^50 51^) and supra basal (Cld4, Dsg1a, Cdkn1a (P21)^52 53 54^) cells indicated that cluster 1 represents a population of basal conjunctival cells while cluster 2 denotes supra-basal conjunctival cells (Fig. 1g-h). Krt15 was highly expressed by conjunctival basal cells (cluster 1) and even higher in conjunctival supra-basal cells (cluster 2), whereas Krt14 displayed basal cell enrichment in conjunctival cells (Fig. 1i).

#### Limbal cells

Clusters 3-5 were identified as limbal cells based on the lack or lower expression of conjunctival (Fig. 1c, S1c) and corneal (Fig. 1d) markers. While cluster 5 displayed supra-basal cell phenotype, cluster 3-4 were identified as limbal basal cells (Fig. 1g-h) expressing a set of new limbal specific markers (Fig. 1e-f). As compared to cluster 4, cluster 3 was hallmarked by much higher levels of Krt15 and Krt14 (Fig. 1i) and lower Krt12 (Fig. 1d), suggesting that it represents the most primitive undifferentiated limbal cell population.

#### Corneal cells

Although a minimal basal expression levels of corneal specific Krt12 mRNA was found in all groups, clusters 6-9 expressed much higher levels of Krt12 (Krt12^hi^) (Fig. 1d). Clusters 6-7 were basal corneal epithelial markers although cluster 7 appeared to include cells that seemingly initiated differentiation, as evident by attenuation of basal epithelial cell markers on the expense of supra-basal cell markers (Fig. 1g-h). Likewise, analysis of basal/supra-basal cell markers suggested that cluster 8 represents partially differentiated cells, most likely corneal wing cells, whereas cluster 9 represents terminally differentiated superficial cells (Fig. 1g-h). In line with previous reports^33 41^, while Krt15 and Krt15-GFP labeled basal conjunctival/limbal epithelial cells, they also marked supra-basal cells of the conjunctiva, limbus and marginal cornea periphery (Fig. 1i). Interestingly, the signal of endogenous Krt15 did not overlap with that of Krt15-GFP (Fig. 1l), in line with previous reports on epidermis^55^. This suggests that the genomic insertion site of K15-GFP transgene, the fractional Krt15 promoter cloned, lack of Krt15 coding sequence or presence of GFP sequence, may have deviated the transcriptional regulation of the Krt15-GFP transgene away from that of endogenous Krt15.

#### Cells in mitosis

Interestingly, clusters 10-11 were found to represent cells from all 3 tissues (conjunctiva, limbus, cornea (Figs. 1c-f)) that were hallmarked by various cell cycle genes including Mki67, Top2a, Ccna2 (Cyclin A2) (Fig. 1j). The grouping of this cell population of mixed lineages into discrete clusters indicated that the changes in expression of genes related to mitosis were profound and dominated the differential expression of tissue-specific genes. Further analysis of cell cycle genes, suggested that cluster 10 represents cells that were captured at early stage of mitosis (late G1-S) while cells of cluster 11 were netted at relatively more advanced stages (S, G2 and M) of the cell cycle (Fig. S1d).

#### Two novel limbal sub-compartments

To certify the cluster identification *in vivo*, we performed in situ hybridization (ISH) on tissue sections to assess the anatomical localization of selected clusters. Eyes of 2-3 months old C57BL/6 mice were enucleated and tissue sections were stained as detailed in Methods. The conjunctiva and cornea were labeled using ISH probes for Keratin 4 (Krt4) and Krt12, respectively. In order to identify the location of the limbal clusters 3-4, we focused on Gpha2 and Atf3 that displayed relatively high mRNA copies. Interestingly, this analysis revealed 2 novel well-demarcated sub-compartments in the limbus. The “outer limbus” was occupied by thin epithelium comprised of small basal epithelial cells with flattened morphology that expressed Gpha2 mRNA. The basal layer cells of the “inner limbus” expressed Atf3 mRNA and displayed flattened cell morphology that gradually became cuboidal towards the corneal periphery (Fig. 1k). In agreement, immunostaining showed that the outer limbal basal epithelial cells were Krt15^+^/Ifitm3^+^/Cd63^+^ whereas inner limbal epithelial cells were Krt15-GFP^+^/Atf3^+^/Mt1-2^+^ (Fig. 1l).

It was noticeable that the two limbal sub-compartments are thin. Moreover, the outer limbus tissue was occasionally absent or truncated in tissue sections. Therefore, to keep the tissue intact and facilitate efficient analysis, we established wholemount immunostaining for antibodies that were found suitable. Figure 1m-o, shows that antibodies against Gpha2, Ifitm3 and Cd63 specifically labeled the outer limbus while Krt15-GFP and Mt1 labeled the inner limbus by wholemount immunostaining.

For consistency and further analyses, we defined conventions for the identification of the outer and inner limbus. The outer limbus markers (Gpha2, Ifitm3 or Cd63) displayed uniform and sharp signal and were well-correlated with slow proliferation and clonal growth dynamics (Figs 2-3). Hence, these markers hereafter were chosen to define the outer limbus, as illustrated in Fig. 1o. Next, the inner limbus was defined as the limbal zone that was negative to outer limbal markers, and its boundary with the peripheral cornea was demarcated by inner limbus markers (e.g. Krt15-GFP in Fig. 1o). Accordingly, quantitative analysis showed that the average width of the outer limbus was ∼100μm while the inner limbus was much wider (∼240μm) (Fig. 1p). Interestingly, the pattern of Krt15-GFP signal was often continuous, or nearly continuous with outer limbus markers like Gpha2, but never overlapped with the labeling of the outer limbus (Fig. 1n-o). Interestingly, by microscopic analysis of wholemount tissues, it was noticed that flattened basal cells found in both sub-compartments were hallmarked by a unique pattern of nuclei staining that differed from that of the cuboidal basal cells that were detected in the marginal inner limbus and corneal periphery (Fig. S1e-f). Finally, analysis of selected keratin coding genes confirmed that basal outer limbal epithelial cells were Krt15^hi^/Krt14^hi^/Krt17^med^/Krt12^neg^, basal inner limbal epithelial cells were Krt15^neg^/Krt14^med^/Krt17^neg^/Krt12^low^, while conjunctival cells were Krt19^hi^/Krt17^hi^/Krt6^hi^/Krt8^med^/Krt12^neg^ (Fig. S1c, g-h).

**Fig 2:**
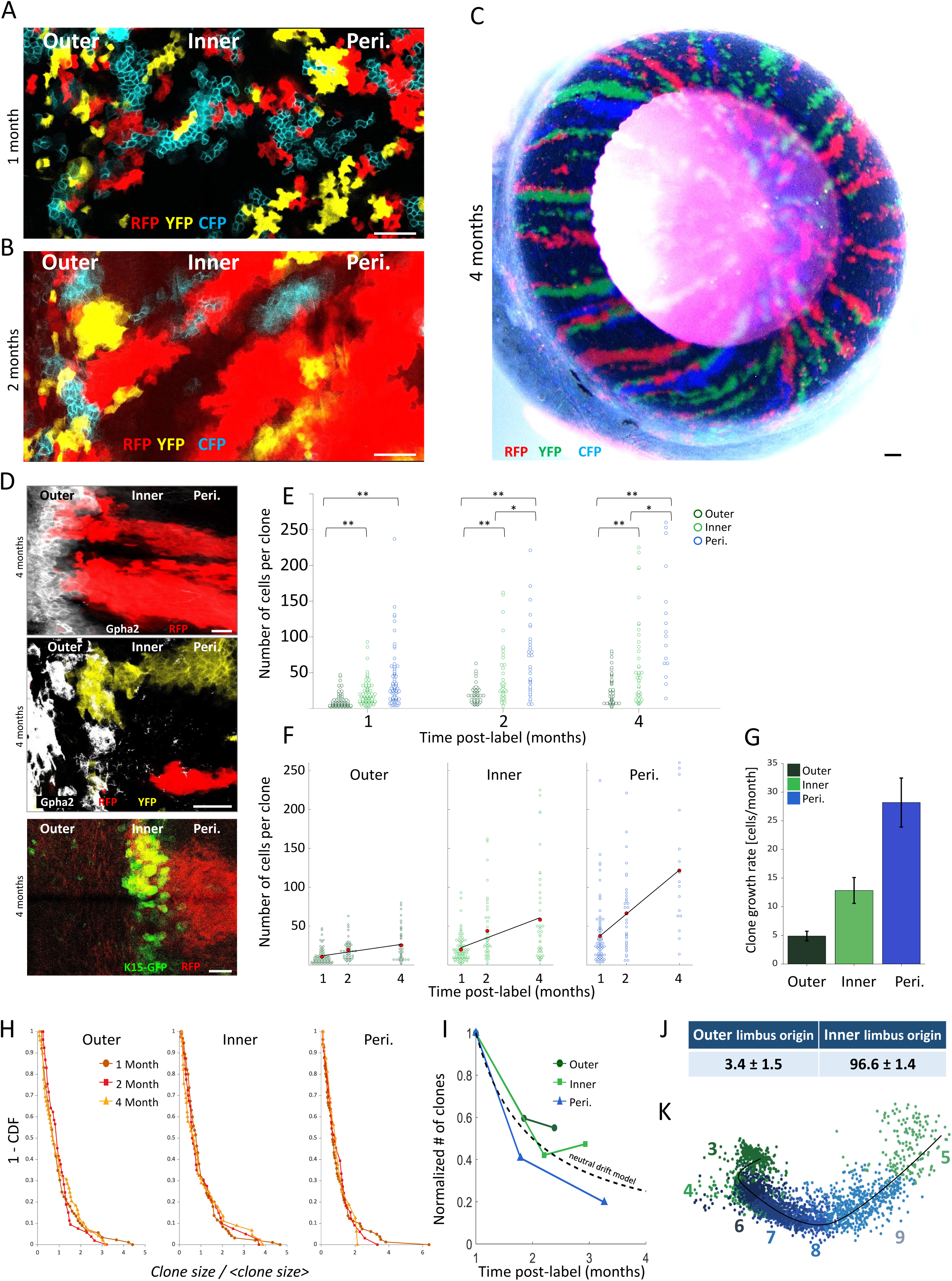
Quantitative “Confetti” lineage tracing unravels the dynamics of outer and inner LSCs. Lineage tracing was induced by tamoxifen injection in 2-month-old UBC-Cre; Brainbow2.1 mice. Confetti-positive clones were analyzed at the indicated time points post-induction. Typical confocal images of the limbal zones are shown in (A-B) and (D), and an entire cornea is shown in (C). The same pictures shown here in A-B, are also shown together with Gpha2 staining in Fig. S4D-E. (E) Scatter plot of the clone size distributions for the outer, inner, and periphery regions at different time points. (F) Clone size evolution in time for the different colonial regions. Red dots denote the average clone size and the black line indicates the linear model that fits the data. (G) The clonal effective growth rate, as estimated from the linear fit. Error bars are 95% confidence interval (CI). Note that the 95% CI does not overlap. (H) One minus the cumulative distribution function of the clone size divided by the mean clone size. The scaled distributions collapse onto the same curve (compare to Fig. S4G, which shows the non-scaled distributions). (I) The normalized number of clones as a function of different time points in different regions. The dotted line denotes the inverse relation between clone size and clone number as expected from the neutral drift model. (J) The percentage of stripes that emerged from the outer or inner limbus. (K) PCA plot predicts the differentiation process across clusters. Scale bars are 100 μm and 50 μm for (C). Statistical significance was calculated using the Kolmogorov-Smirnov test. (*, p-value < 0.05; ** p-value <0.005). Abbreviations: Conj., Conjunctiva; Peri., Periphery.

#### In silico analysis of clusters

Next, pathway enrichment analysis using WebGestalt^56^ was performed with the ORA method using KEGG as the functional database in order to identify potential pathways that were enriched in clusters 3 or 4 in comparison to all other clusters. Interestingly, SC and cancer pathways including Pi3K and P53 were enriched in cluster 3 while cluster 4 was enriched with TNF pathways and multiple annotations for cancer (Fig. S2). Additional analysis revealed a list of differentially expressed genes expressed by clusters 3 or 4 (Fig. S3a-b). Expectedly, genes related to hemi-desmosomes increased in clusters of basal cells (clusters 1, 3-4, 6-7), and desmosome-related and gap junction associated genes increased in clustered supra-basal cells (clusters 2, 5, 8-9) (Fig. S3c). Interestingly, high levels of the extracellular matrix (ECM) components Lamb3 and Crim1 (Cysteine-Rich Motor Neuron 1), that regulates BMP and inhibits cell proliferation^57^, were found in the outer limbus cluster 3, whereas Col17a1 was higher in the inner limbus cluster 4. Taken together, this data illuminates the genetic signature of diverse cell states across the corneal epithelial lineage, and in particular, reveals the co-existence of two discrete limbal sub-compartments consisting of two distinct limbal epithelial cell populations.

### Quantitative lineage tracing suggested distinct clonal dynamics and lineage hierarchy for outer and inner limbal epithelial cells

To further uncover the nature of outer and inner limbal epithelial cells, functional enrichment analysis (WebGestalt with ORA and KEGG) shed light on differences between clusters 3 and 4. Interestingly, among the top 10 functional categories, 4 identifiers involved cell proliferation while 3 involved cell movement (Fig. S4a), suggesting that a central difference between the limbal clusters 3 and 4 lies in their cell cycle and motility characteristics. We thus aimed to gain insights regarding proliferation, clonal dynamics, centripetal movement, and discover the potential hierarchy between all clusters and within each limbal zone. For that purpose, we designed a quantitative genetic lineage tracing experiment for tracking all cell populations including potentially rare LSC populations, in a clonal and inducible manner. We established double transgenic UBC-Cre^ERT2^; Brainbow^2.1^ mice that facilitated the tamoxifen inducible activation of Cre recombinase in all cell types in an efficient and unbiased manner under the control of the ubiquitin C (UBC) promoter (Fig. S4b). In the absence of tamoxifen, Cre is expressed at the cytoplasm of all cell types of this mouse, thus unable to access the DNA. However, upon transient exposure to tamoxifen, Cre recombinase translocates into the nucleus where it can randomly induce DNA excision and/or inversion of the loxP sites in the brainbow^2.1^ cassette, thus, allowing stochastic and irreversible expression of one out of four “confetti” fluorescent proteins (namely, nuclear green (GFP), cytoplasmic red (RFP), cytoplasmic yellow (YFP) or membrane cyan (CFP) fluorescent protein) (Fig. S4c)^58^.

Two-month-old mice were treated with tamoxifen for 3-4 consecutive days to induce efficient “Confetti” labeling in all cell populations. For quantitative analysis of the limbal sub-compartments, wholemount staining of Gpha2 was performed to identify the outer limbus (647nm to avoid overlap with 4 Confetti colors and DAPI) while the contiguous 100μm Gpha2-negative limbus was considered as inner limbus for calculations (Fig. S4d).

Interestingly, at all time-points, the distribution of clones size (i.e., the number of basal cells in a clone) in the outer limbal epithelial zone was significantly lower compared to those found in the inner limbus and the peripheral cornea. Moreover, the average clonal size in the inner limbus was smaller than the size of clones in the corneal periphery (Fig. 2a-d, e). Two months post induction, a pattern of radial stripes emerged already in clones that held a “foot” in the inner limbus (Fig. 2b). Four-months post induction, when the pattern of fully developed, radial stripes appeared. Most stripes extended from the inner limbus and only a few confetti stripes were sourced from the outer limbus (Fig. 2d, j). The detection of stripes starting from both outer and inner limbal zones after very long-term tracing, suggests that both limbal zones contain LSCs.

The uniform expression pattern of markers and clonal growth dynamics implied that the outer limbal epithelial cells might follow stochastic rules and neutral drift^59^. A main prediction of this approach is that the size of the clone would grow linearly with time^60^.In this case, the slope of this linear trend is proportional to the product of doubling rate and the probability of symmetric division in the epithelial plane (Methods). Our results suggest that clone size dynamics in all regimes can be captured by a linear growth and that the slopes are significantly different and ordered centripetally– each regime has a slope which is about twice higher than its neighboring zone (Fig. 2f-g). Another prediction of the neutral drift model, is that the clonal size distributions scale with their averages. That is, the size distribution of the clone size normalized by their mean should be similar. Fig. 2h shows the probability of having a clone that is larger than some value (one minus the cumulative probability distribution) for the normalized size distributions. All regimes show clear scaling of their clone size for all time points. Furthermore, the neutral drift model predicts that the number of clones should decrease inversely with the clone size (Fig. 2i). In our case, there is a clear difference between the three different zones tested. While in the periphery the expected inverse relationship between size and number is similar to that of neutral drift, in the outer and inner regimes after four months the reduction in the number of clones is smaller than expected by that model. Taken together, this data suggests that the outer limbus and inner limbus contain a population of SCs that have similar stochastic dynamics but different doubling times.

If the outer LSCs are the most primitive undifferentiated cells positioned at the tip of the cellular hierarchy, the inner limbus may host slightly committed LSCs. To further assess the hierarchical relations between clusters in the corneal epithelial lineage, we plotted the clusters using linear dimensional reduction using the Seurat algorithm for limbal/corneal epithelial clusters (clusters 3-9). This algorithm infers the sequential changes of gene expression between clusters and thereby provides insights for the dynamics of biological processes, in this case cell differentiation. In line with lineage tracing data, this analysis suggested that cluster 3 (outer limbal basal cells) gives rise to cluster 4 (inner limbal basal cells) that derive clusters 6-7 (corneal basal cells) and clusters 8-9 (corneal wing and superficial cells), while cluster 5 cells (limbal superficial) that are indeed very similar to corneal superficial cells were positioned last (Fig. 2k). Taken together, the combined quantitative lineage tracing and in silico analysis illuminates the dynamics and hierarchy between corneal epithelial cell populations.

### Outer and inner limbus represent segregated sub-compartments for quiescent and activated LSCs

To rigorously test LSC growth dynamics by an alternative and direct approach, we next examined the cycling properties of outer and inner basal limbal epithelial cells by staining of Ki67 which labels cells that are at different stages of cell division (late G1, S, G2, M). In agreement, only 5.7±3.0% of outer limbal basal cells were Ki67^+^, while 16.5±3.9% of inner limbal basal cells and 19.8±4.0% of corneal peripheral basal cells were Ki67^+^ (Fig. 3a-b). To corroborate this data, we performed 5-Ethynyl-2’-deoxyuridine (EdU) nucleotide analogue incorporation assay to identify undergoing DNA replication (S-phase). As illustrated in Fig. 3c, 2-month-old mice were intra-peritoneal injected with EdU, six hours later mice were sacrificed and cells that were in the S-phase of cell division during the interval were identified as EdU^+^ cells by whole mount staining. Expectedly, EdU incorporation by outer limbal cells was much less frequent, as compared to that of basal epithelial cells at the inner limbus or peripheral cornea (Fig. 3d). To examine the ability of outer limbal cells to enter active cell cycle in response to corneal injury, we performed large (2.5mm) corneal epithelial debridement following EdU injection. Evidently, 24-hours post injury, numerous outer (and inner) limbal cells entered cell cycle and were EdU positive. Taken together, this set of experiments suggests that the outer LSCs are slow-cycling cells, but they can quickly enter cell cycle and participate in corneal healing. To further examine the presence of slow cycling cells in the outer limbus, we performed an EdU pulse-chase experiment. As described in Fig. 3e, mice were injected with EdU twice a day for 15-days (“pulse”) followed by a “chase” period of 30-days at the absence of EdU. During the chase period the label declines by 50% within every cell division and the label becomes undetectable after few successive division rounds (∼4), depending on assay type and sensitivity^61 38^. As expected, EdU^+^ label retaining cells were predominantly found in the outer limbus, whereas rare cells were occasionally found in the inner limbus (Fig. 3f), suggesting that during the 30-day chase period most quiescent outer LSCs exceeded sufficient division rounds and lost labeling and only few which underwent fewer replications could be detected. To validate that label retaining cells were epithelial cells, we performed co-staining of EdU^+^ label retaining cells with Krt17 antibody (Fig. S1h).

**Fig 3:**
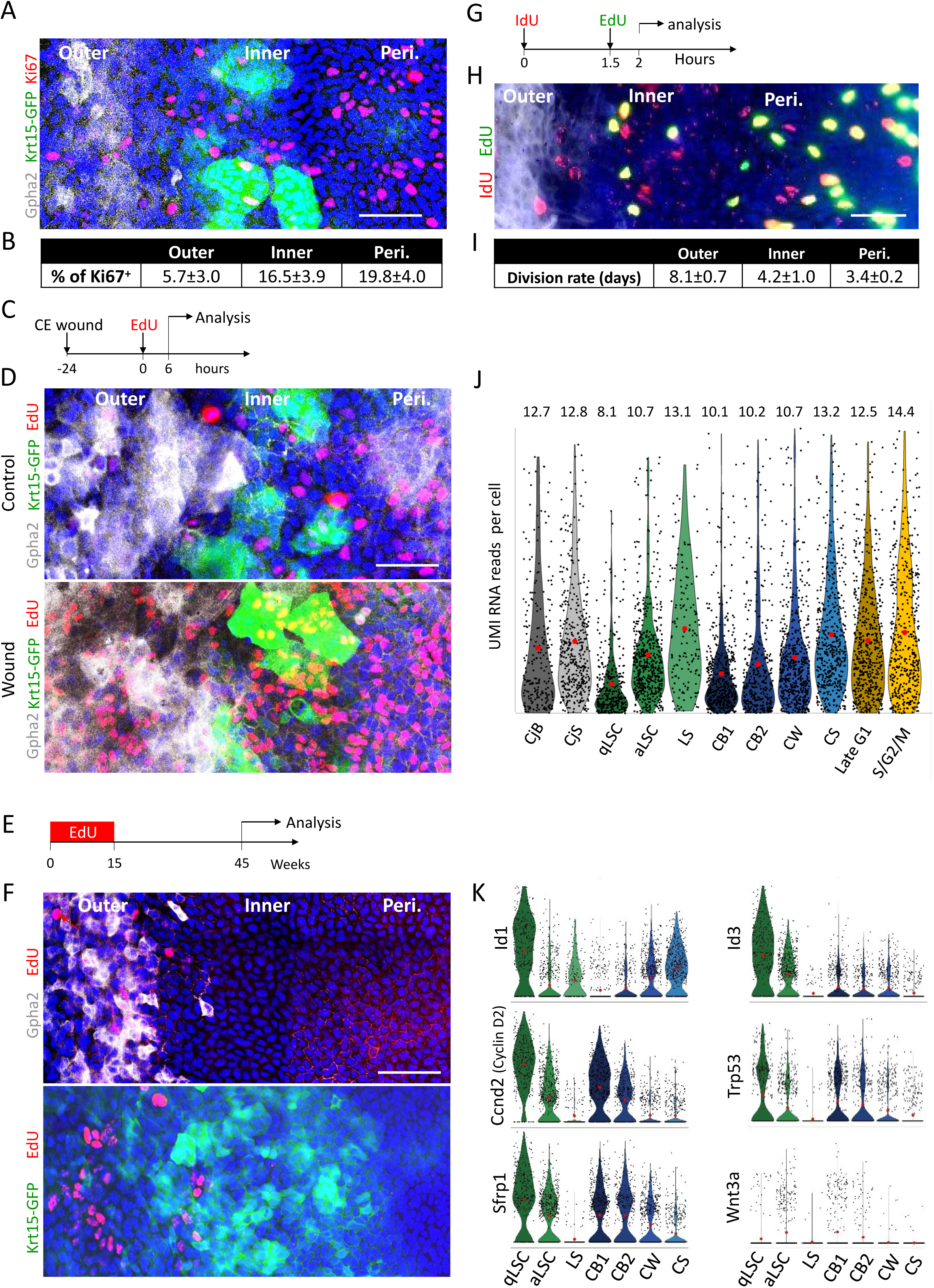
Proliferation analysis of slow cycling and frequently dividing LSCs. Adult 2-3 month old mice were used for all experiments. The limbal region markers in wholemount corneas were defined by Gpha2/K15-GFP, nuclei were detected by DAPI counter staining (A, D, F, H) while Ki67^+^ cells were stained (A) and quantified (B). Schematic illustration of the wound experiment (C). Corneal epithelial (CE) wound was performed and 24-hours later, EdU was injected and 6 hours later, cells in S-phase were identified (EdU^+^) in wounded or uninjured controls. Typical confocal image of whole mount immunostaining shown in (D). (E) Schematic illustration of pulse-chase experiments performed to identify slow cycling EdU^+^ label retaining cells, and typical confocal image of whole mount staining is shown in (F). (G) Schematic illustration of double nucleotide labeling experiments performed to estimate division frequency of cell populations and typical confocal image of whole mount staining is shown in (H), and calculated estimation of division rates are shown in (I). (J) Violin plot diagram of unique molecular identifiers (UMI) RNA reads per cell in each cluster. (K) Violin plot showing the expression of the indicated genes. Statistical significance was calculated using One-way Anova test followed by Bonferroni test (*, p-value<0.05). Data represent mean ± standard deviation, Abbreviations: Peri., Periphery; CjB, Conjunctival Basal; CjS, Conjunctival Suprabasal; qLSC, quiescent limbal stem cell; aLSC, activated limbal stem cell; LS, limbal superficial; CB1, corneal basal 1; CB2, corneal basal 2; CW, corneal wing; CS, corneal superficial. Scale bars, 50 μm.

The relative homogeneity in the expression of Gpha2/Ifitm3/Cd63 by the outer LSCs, together with the absence of Ki67 associated with broad slow growth of clones in the outer limbus, implied that the vast majority, perhaps entire, basal cell population in the outer limbal zone divides infrequently. To gain direct evidence on cell cycle of each population and evaluate the rate of cell division in each zone we performed a double nucleotide-analogue injection. As depicted in Fig. 3g, 1.5 hours following injection of iododeoxyuridine (IdU, red), EdU (green) was injected, and 30 minutes later tissues were harvested, stained and analyzed (Fig. 3h-i). As detailed in Methods, S-phase length was first calculated as the ratio between the number of cells that were IdU^+^/EdU^+^ (cells that were engaged with, but have not completed, DNA replication) and the number of cells that were IdU^+^/EdU^-^ (cells that were in S-phase after IdU injection but have completed DNA replication already before EdU injection), multiplied by the interval length (1.5 hours). The estimated division frequency was calculated as the ratio between the total number of cells divided by the number of cells in S-phase, multiplied by the S-phase length^62^. This analysis suggests that outer limbal basal cells divide on average every ∼8 days, inner limbal basal cells divide every ∼4 days while corneal peripheral basal cells divide every ∼3.5 days (Fig. 3i). Taken together, this data strongly suggests that the outer limbus contains a predominant equipotent Krt15^+^/Gpha2^+^/Ifitm3^+^/Cd63^+^ cell population of qLSCs while the inner limbus is occupied by a widespreas equipotent aLSCs.

Cellular quiescence is linked with reduced biochemical activity including transcription activity ^16 17^. Indeed, the cells in cluster 3 (qLSCs) displayed the lowest value of RNA reads per cell (Fig. 3j). Previously, BMP was linked with epithelial SC quiescence while WNT repressed BMP pathway and induced SC activation^63 64 65 66^. In agreement, cells of cluster 3 (qLSCs) preferentially expressed the BMP-regulated transcription factors Id1/Id3 and the Wnt inhibitor, Sfrp1^67^, while Wnt3a was higher in cluster 4 of aLSCs (Fig. 3k). Interestingly, cluster 3 expressed Cyclin D2^68 69^and the tumor suppressor Trp53^70^, both of which were linked with growth arrest induced by various stimuli.

### IFITM3 and GPHA2 marked KRT15^+^ human LSCs and IFITM3 supported undifferentiated state

The relevance of qLSC to cell therapy is significant, therefore, we next explored the usefulness of the newly identified markers of murine qLSCs to human. Immunofluorescent staining revealed that KRT15, IFITM3 and GPHA2 are expressed by basal limbal epithelial cells (Fig. 4a). Similar to staining of murine epithelium (Fig. 1-3), the labeling pattern of qLSC markers was not sporadic but rather wide, marking a large population of basal limbal epithelial cells. Interestingly, however, GPHA2 expression was drastically down regulated to hardly detectable levels upon cultivation of LSCs. Repression of GPHA2 (esiGPHA2) resulted in total inhibition of GPHA2 expression (Fig 4b). This implies that GPHA2 expression depends on niche specific signal that was absent *in vitro*, and that GPHA2 is tightly associated with a slow cycling feature that is not maintained *in vitro*. Nevertheless, KRT15 and IFITM3 were highly expressed by undifferentiated limbal epithelial cells *in vivo* and *in vitro* (Fig. 4c). In addition, calcium induced differentiation enhanced KRT12 and reduced KRT15, IFITM3 and P63 (Fig. 4c). In a previous report, IFITM3 was linked with localization to endo-membranes where it protected embryonic SCs from viral infection^71^. In line, IFITM3 signal was confined to cellular vesicle structures in the cytoplasm of cultivated limbal cells (Fig. 4d).

To test whether IFITM3 influences stemness and differentiation, we performed a knock down experiment. To achieve efficient and specific knockdown, we used endoribonuclease-prepared silencing RNA (esiRNA) where enzymatic digestion of long double stranded RNAs produces a mixture of multiple short fragments of silencing sequences, thereby, enhancing specificity and efficiency. Efficient knock down of IFITM3 was evident by RNA and protein levels (Fig. 4e-f). Moreover, repression of IFITM3 resulted in a significant reduction in KRT15, and an increase in the expression of the differentiation associated corneal keratin KRT12 (Fig. 4e-f). Taken together, this set of experiments suggests that IFITM3 positively controls the undifferentiated state of human qLSCs, and that the newly identified qLSC markers (IFITM3, GPHA2) can be used to identify human qLSCs.

**Figure 4:**
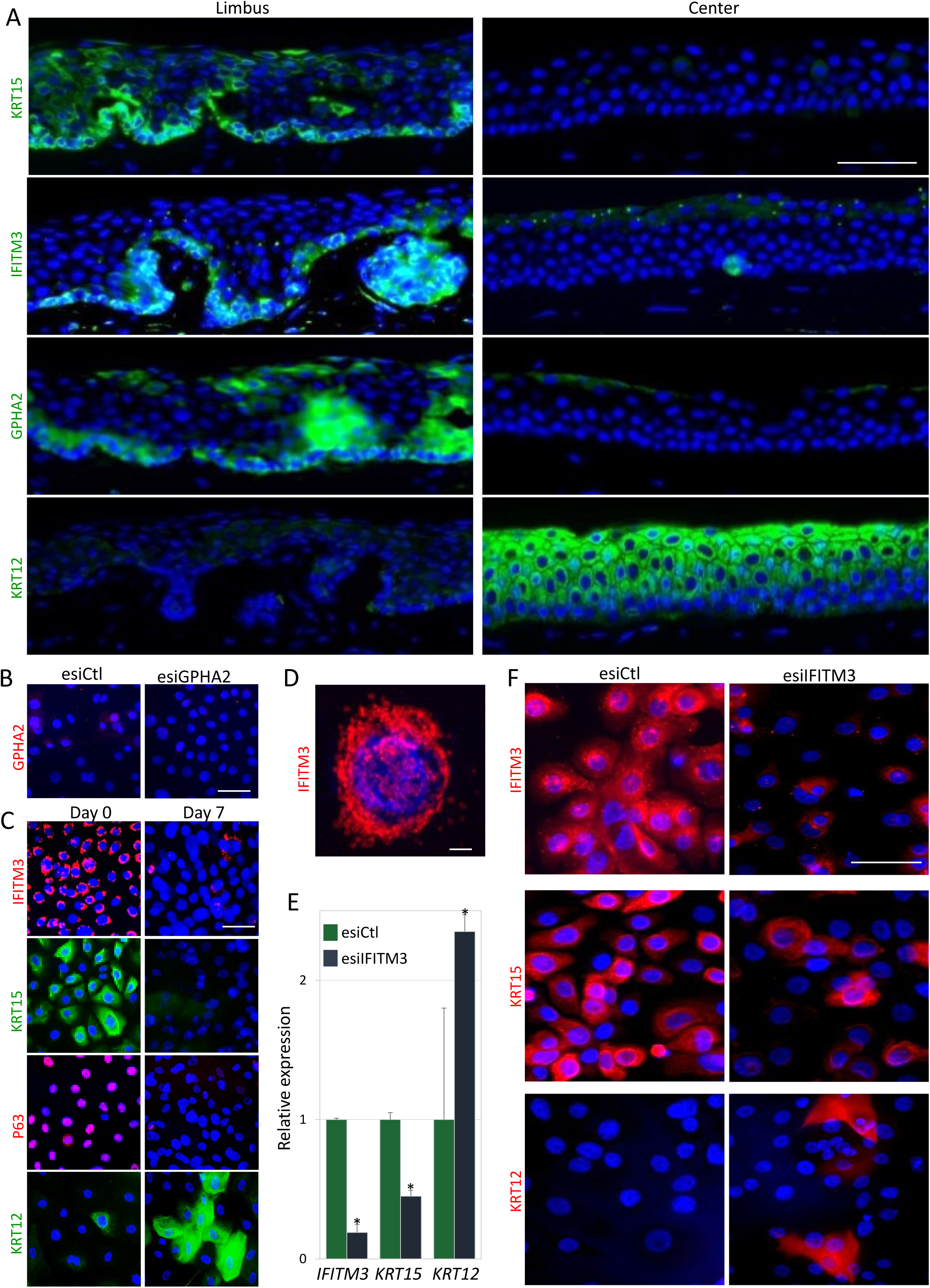
GPHA2 and IFITM3 marked qLSCs while IFITM3 supported undifferentiated state *in vitro*. (A) Human cornea frozen sections were subjected to immunofluorescence staining of indicated markers. (B) Human Limbal epithelial cells were transfected with non-specific control esiCtl and endoribonuclase prepared small interfering RNA against GPHA2 (esiGPHA2), cells were immunostained for GPHA2. (C) Human limbal epithelial cells were cultured and maintained at low calcium (Day 0) or induced to differentiate (Day 7) and stained for the indicated markers. (D) Enlargement of IFITM3 staining shown in (C) (day 0). (E-F) Undifferentiated cells were transfected with esIFITM3 or esiCtl. The expression of the indicated markers was examined 4-5 days later, by quantitative real-time polymerase chain reaction (E) and immunofluerscent staining (F). Nuclei were detected by DAPI counter staining (A-D, F). Scale bars are 50 μm or 10 μm (D). Statistical significance was calculated using t-test (*, p-value<0.05).

### T cells serve as niche cells for qLSC regulating cell proliferation and wound closure

The striking segregation of qLSCs to the outer limbus suggests that this cell state must be regulated by the defined local microenvironment (niche). Interestingly, the outer limbus was clearly demarcated by well-structured blood/lymph vasculature (Fig 5a). Thin well-organized blood vessels typically penetrated the marginal inner limbus but never reached the corneal periphery. Lymph vessels were characteristically present in the outer but not the inner limbus. Additionally, the limbus was highly populated with Cd45^+^ immune cells, amongst them, many Cd45^+^ that displayed dendritic cell morphology, both in the stroma and in the epithelium (Fig. S5a). Interestingly, we have identified a population of T cells in the outer limbus, many of which expressed the regulatory T cell markers, Cd4 and Foxp3 (Fig. 5b, S5a) and others expressed the cytotoxic T cell marker Cd8 (Fig. S5b), suggesting that qLSCs may be regulated by T cells.

Next, we explored the limbus of two laboratory mouse strains that fail to maturate T and B lymphocytes, namely, severe combined immunodeficiency (SCID) and non-obese diabetic SCID (NOD/SCID), and BALB/c mice served as a control group. Similar to C57BL/6 genetic background (Fig 1-3), the outer limbus of BALB/c mice was hallmarked by Gpha2^+^/Cd63^+^/Ifitm3^+^ that infrequently expressed Ki67 (Fig. 5b-d, S5a-c, S5e). Curiously, an extensive reduction of Gpha2 and Cd63 proteins to levels that became barely detectable was found in SCID mice (Fig. 5b-c) as well as in NOD/SCID mice (Fig. S5c), while other qLSC markers (e.g. Ifitm3) were unaffected (Fig. S5b). Moreover, higher index of Ki67 labeling was found in the outer limbus (as well as inner limbus) of SCID mice (Fig. 5c-d) that displayed mild epithelial thickening (Fig. S5d). In agreement, similar defects were recapitulated in NOD/SCID mice (Fig. S5c). Since no B cells (Cd19^+^) were detected in the limbus (Fig. S5b), we assumed that this phenotype is caused by the absence of T cells in SCID mice. To further address the involvement of T cells in qLSC regulation, we explored athymic Nude-Foxn1nu mice that lack T cell development. Here too, qLSC markers Gpha2 and Cd63 were dramatically attenuated while Ifitm3 was maintained, strengthening the involvement of T cells in qLSC regulation (Fig. S5e). Finally, we aimed to substantiate the crosstalk of T cells – qLSC in adulthood, and exclude the possibility of developmental failure. To this end, we repressed the ocular immune system by topically applying the corticosteroid Dexamethasone (Fig. S5f), or alternatively and more specifically, performed inhibition of regulatory T cells by sub-conjunctival injection of anti-Cd25 antibody (PC61.5). Intriguingly, 6-days post antibody injection, Cd63 and Gpha2 drastically decreased whereas cell proliferation increased (Fig. 5e-f), strongly suggesting that T cells directly regulate quiescence in the outer limbus. Similar results were obtained following Dexamethasone treatment (Fig. S5f). Finally, to link between T cell regulation and qLSC functionality, we performed corneal epithelial debridement and followed epithelial closure by fluorescein dye penetration. As shown in Fig. 5g-h, immunodeficient mice displayed delayed wound closure. Taken together, this set of experiments suggests that T cells, most likely Cd4+/Foxp3+ regulatory T cells, serve as a niche for qLSCs and play a critical role in quiescence maintenance, control of epithelial thickness and wound healing.

**Figure 5:**
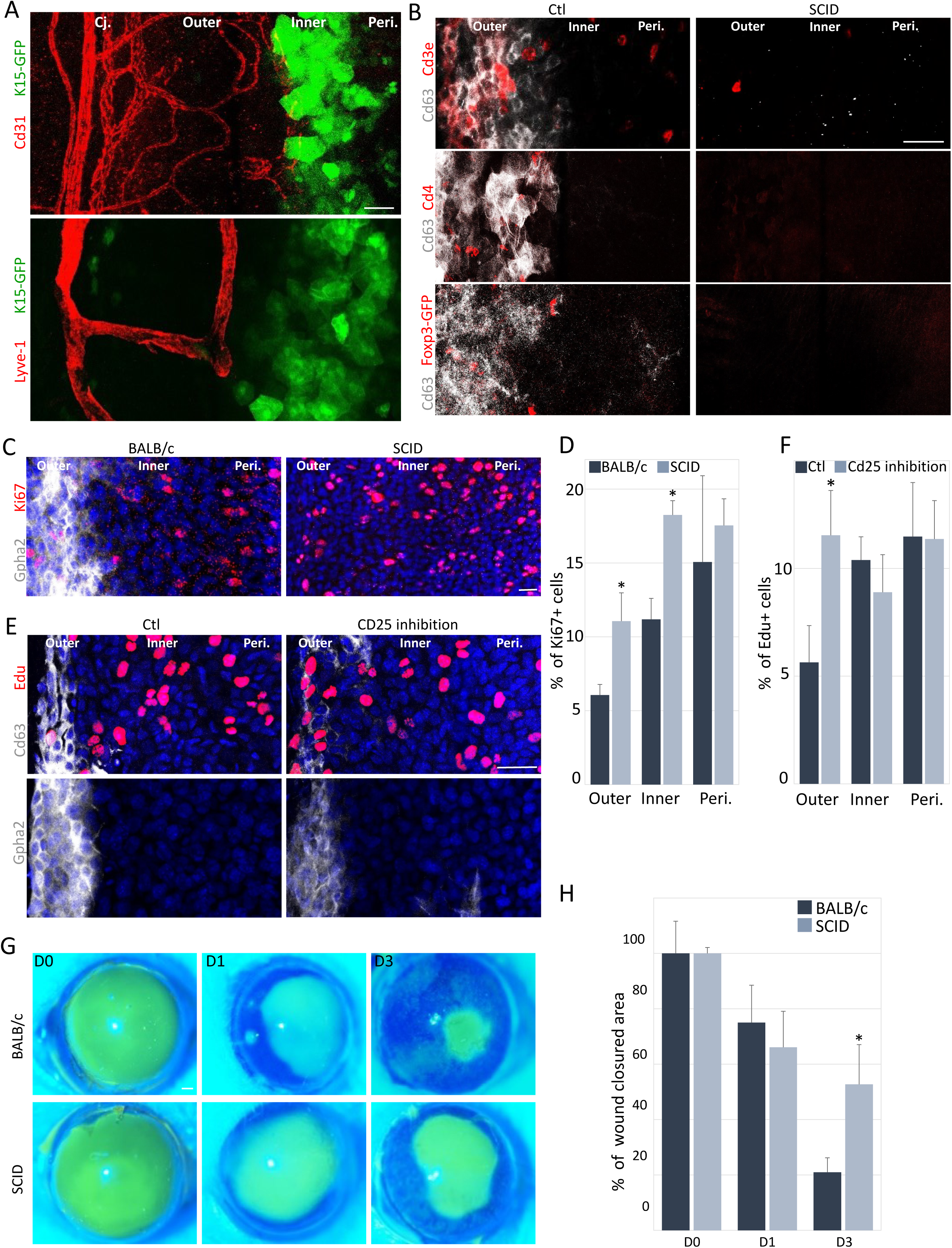
T cells regulate qLSC proliferation and response to wound stimulus. (A) Whole-mount immunostaining of corneas was performed to label blood (Cd31) and lymph (Lyve-1) vessels of 2-3 months old Krt15-GFP mice. (B-C) Whole-mount immunostaining of corneas of immunodeficient mice (SCID) or controls to identify qLSCs (Cd63^+^ or Gpha2^+^) or T cell populations (Cde3^+^ and Cd4^+^, or FoxP3-GFP^+^ transgene) or dividing cells (Ki67^+^). (D) Quantification of Ki67 expression in different limbal zones. (E) Six-days post subconjunctival injection of anti-Cd25 antibody (inhibits regulatory T cells) to Krt15-GFP mice, whole mount immunostaining of the indicated qLSC markers was performed. (F) Quantification of EdU expression in different limbal zones. (G-H) Corneal epithelial debridement was performed in 2-month old mice and fluorescein dye stain was performed to follow wound closure. Typical images are shown in (G) and quantification in (H). Nuclei were detected by DAPI counter staining (C, E). Scale bars are 50 μm. Statistical significance was calculated using t-test (*, p-value<0.05). Abbreviations: Cj., Conjunctiva; Peri., Periphery.

## Discussion

In 1989, Cotsarelis and Lavker reported the identification of rare slow cycling cells in the limbus^10^. Since then, over the last 3-decades, much attention has been given for identifying such rare cells, however, no reliable marker has been found yet. Here we captured the signature of primitive qLSCs and propose that these cells are more prevalent than previously estimated. We showed evidence that qLSCs, which populate the basal layer of the outer limbus, quite uniformly express Krt15^+^/Gpha2^+^/Ifitm3^+^/Cd63^+^. The ∼8 days estimated cell cycle length of qLSC fits well with quantitative lineage tracing, double nucleotide incorporation experiments, and nucleotide pulse-chase label retention assays.

Indeed, the limbal/corneal epithelium becomes uniformly labeled following 2-weeks of nucleotide pulse^38 10^, i.e., suggesting that the length of cell cycle of both limbal and corneal epithelial cells is shorter than 14 days. Additionally, if assuming ∼8 days for qLSCs, the interval of 30-days of chase in the absence of EdU (Fig. 3e-f) should theoretically allow ∼4 division rounds for outer qLSCs, and it is therefore not surprising that most outer limbal basal cells lost labeling while only few retained low EdU signal. In other words, this data fits well with abundant qLSCs model. Likewise, this model is supported by the quantitative analysis of Ki67 staining and EdU incorporation after short pulse (Fig. 3a-d) as in both cases, the outer limbus showed a widespread negativity for labeling. Our working model that posits abundancy of equipotent qLSCs fits well with a stochastic SC model that was proposed to describe SC dynamics in the epidermis, esophagus and gut epithelium^23 58 72^. In these tissues, abundant equipotent SCs that were shown to be extremely fast cycling, are at the tip of the cellular hierarchy. Epidermal and esophageal SCs divide every 2-3 days while gut epithelial SCs divide once a day, much faster as compared to qLSCs that divide every 8 days. This suggests that under distinct physiological conditions, tissues display tailored regenerative strategies for reasons of cost and benefit tradeoff.

The two new LSC populations were well-segregated into well-defined limbal sub-compartments. This conclusion is supported by the observations that the outer and inner limbal cells differentially express specific markers and engaged pathways (Fig. 1, S2-4), display distinct clonal growth and proliferation dynamics (Fig. 2-3), and niche components (Fig 5). The lineage tracing experiments implies that aLSCs play a key role in cornea replenishment under homeostasis, as most stripes emerged from this region. By contrast, a key feature of qLSCs is their ability to rapidly exit dormant state and enter the cell cycle in response to injury^73^ (Fig. 3).

The cell cycle length of qLSCs is only approximately twice longer than that of aLSCs that were estimated to divide every ∼4 days. However, this frequency of cell division is higher compared to some estimations for other tissue-specific slow cycling SCs^74 75^. The reason for such a drastic difference in SC cycling frequencies and its importance is poorly understood. In this sense, one could not exclude the possibility that an even slower cell population co-exists in the outer limbus, however, if it does, these cells must be very rare. Moreover, sub-fractionation of cluster 3 of qLSCs does not provide statistically relevant sub-clusters, again, suggesting the cells in the cluster were relatively highly homogenous.

The analysis of the clone size distributions showed that they scale with the average number of clones in all the regions. This suggests that the underlying stochastic dynamics of all the regions are similar. The clone size dynamics are consistent with the neutral drift dynamics model and suggest that the doubling time in the inner limbus is twice the doubling time in the outer limbal regime, which is consistent with the cell cycle length measurements. In all regimes, the number of clones is going down with time, as expected. However, analyzing the quantitative relation between clone size and clone number reveals a striking difference between the outer and inner limbal regimes and the periphery. While all zones exhibit clone size distributions that are consistent with the neutral drift model, the decrease in the inner and outer regimes is slower than expected by this model. This saturation in clone number decrease rate might indicate that the clonal neutral competition is limited in these areas. This could be due to the tight spatial boundary conditions of the limbal regimes.

Our working model (Fig. 6) is in agreement with the hematopoietic, hair follicle^18^, neural^20^ and muscle^19^ lineages, all of which incorporate qSCs that can produce aSCs^18^. What are the benefits and costs of having two distinct SC states? What is the mechanism that controls the transition between SC states? Is this model universal or otherwise only applies for particular tissues? Which physiological constraints influence SC division rates? Interestingly, epithelial SCs of the gut proliferate once a day^76^, while epidermal^77^ and esophageal^78^ SCs divide every ∼3 days. In this context, it is tempting to hypothesize that the corneal transparency and location in the front of the eye makes it an extremely hazardous environment for SCs that must be well guarded especially because they must constantly proliferate to replenish corneal center with new cells. This must entail fundamental considerations for SC well-being such as localizing the SCs at the most well-protected zone (outer limbus) and accrediting SCs with infrequent division.

**Figure 6:**
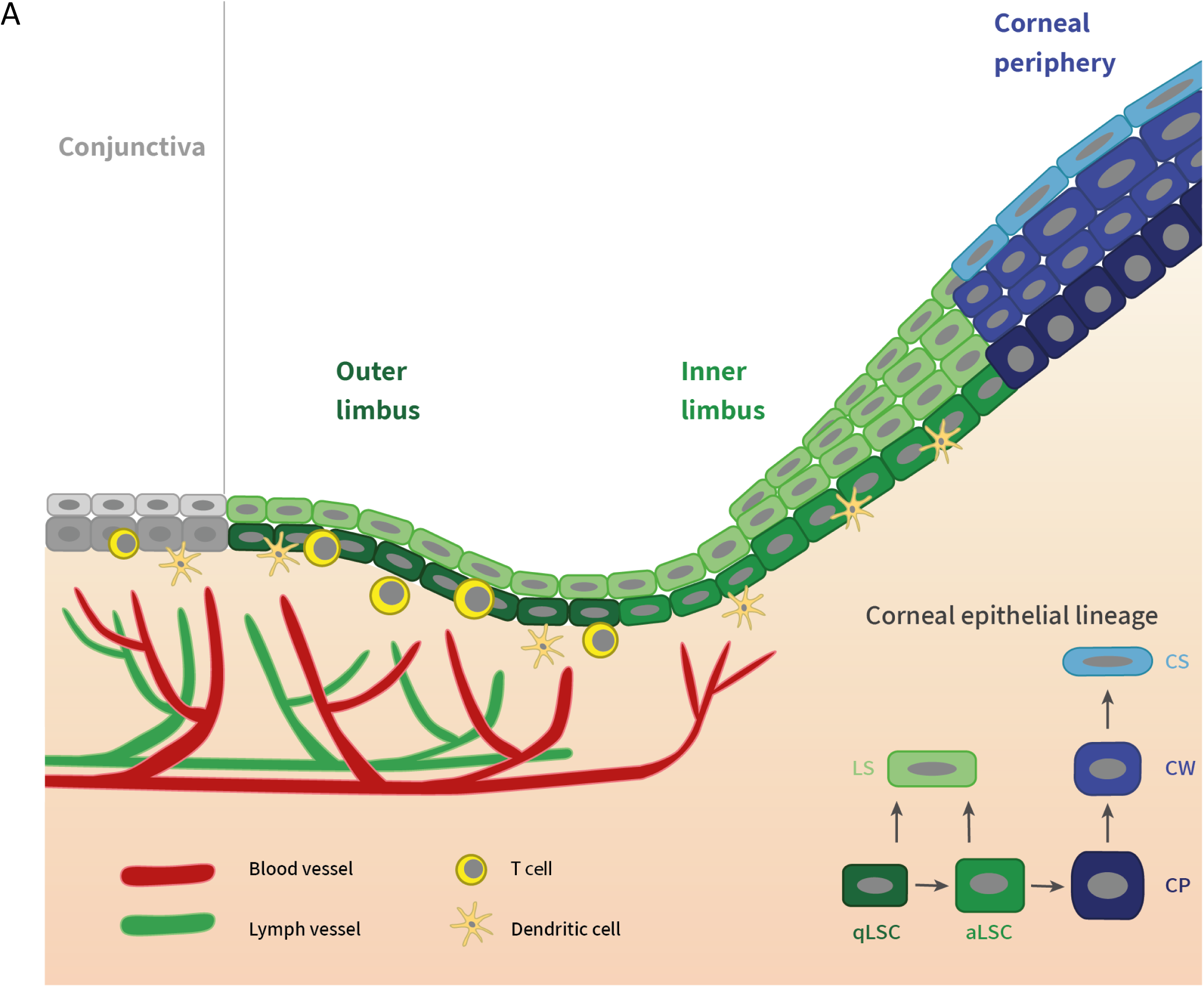
Working hypothesis. Schematic illustration of our working model, depicting the anatomy of the limbal sub-compartments and LSC populations. We propose that the outer limbus is occupied by widespread qLSCs that play an important role as a SC reservoir for wound repair, barrier maintenance and corneal replenishment. The inner limbus contains abundant aLSCs that are actively engaged in corneal replenishment. The maintenance and differentiation of LSC populations is regulated by niche components including blood and lymph vasculature, stromal cells, and particularly T cells that control qLSC proliferation.

The quiescence state of SCs is believed to be accompanied by reduction in DNA replication, metabolism, gene transcription and protein translation^16^. Notably, these processes were associated with molecular damage due to the production of toxic agents or errors in biosynthesis of macromolecules^79 80^. In line, attenuation of these activities by SC dormancy was proposed to delay SC aging^17^. Moreover, the reduced cumulating number of cell replications by quiescent SCs, may significantly attenuate the accumulation of mutations. Surprisingly, however, qSCs employ an error prone mechanism of non-homologous end joining to repair their DNA^81 82^. By contrary, frequently dividing SCs engage a non-error prone mechanism of homologous recombination for repairing DNA breaks^83^, and frequent division rate was shown to be required for the activation of repair pathways^84^. In line, frequently dividing SCs were identified in many tissues including the gut^22^, epidermis^85^,and esophagus^24^. These cumulating findings revised the old dogma that considered quiescence as an integral feature of SCs. It also suggested that epithelial SCs are not quiescent and that the high turnover of such tissues is associated by a single pool of active SCs. The present study refines this conclusion and highlight the possibility that qSCs play an important role in corneal epithelial lineage and potentially in other epithelial tissues that may benefit from qSC reservoir.

The regulation of quiescence is only partly understood, however, it clearly involves interactions with niche extracellular matrix proteins, secreted factors, and niche resident cells. At the molecular level, cell cycle inhibitors (e.g. P21, P27 and P57), tumor suppressor genes (e.g. retinoblastoma and P53) and soluble factors (e.g. BMP4, Dkk-1, sFRP, Wif and Shh) were found to positively regulate quiescence, while cytokines and Wnt activators (e.g. noggin) were associated with the exit from quiescence state and the transition into active SC state^18 16^. The bioinformatic analysis of limbal clusters suggests that qLSC engage Pi3K pathway, the BMP-regulated transcription factors Id1/3^63 64 65^, the tumor suppressor gene Trp53^16^, Cyclin D2^68 69^ and the Wnt inhibitor Srf1^86^, all of which were linked with growth arrest. Of interest, aLSCs were enriched for TNF, WNT and Ap-1 pathways (Fig. S2-3) that were linked with proliferation or SC activation. Interestingly, Atf3 which is a structural protein of Ap-1 complex was shown to repress Id1 and enhance epithelial cell proliferation^87^. Future studies will address the mechanism by which these genes are regulated by outer/inner limbus specific ECM factors (Fig. S3), metabolites or vasculature related factors, or influenced by niche cells (e.g. T cells).

Likewise, future experiments will be needed to illuminate the mechanism that controls the quiescence and the transition into active state. The maintenance of LSC specific states, and the transition between them is most likely driven by niche cells (e.g. T cells, keratocytes, vasculature), ECM and/or biomechanical forces. It is therefore not surprising that quiescence (as well as Gpha2 expression) is lost ex vivo, whereas culture conditions are sub-optimal and lacking essential components of the niche. The co-localization of T cells with qLSC strongly implies that these cells may secret cytokines to control LSC quiescence, in line with previous studies where immune cells were identified as niche cells for epithelial SCs^88^. In the clinics, corticosteroids such as dexamethasone are being widely used to prevent uncontrolled corneal inflammation and neo-vascularisation, as these agents are considered as highly effective drugs for nonspecific suppression of inflammation. However, it has been shown that anti-inflammatory steroids must be used for short time as they may have adverse effects including corneal wound healing delay ^89^. The awareness and recognition of such adverse effect is of particular importance given the observed interference of T cells – qLSC crosstalk. Future studies will be needed to illuminate these interactions and translate this knowledge into optimized treatments. Moreover, this knowledge may facilitate optimal qLSC cultivation for curing blindness in patients that suffer from “LSC deficiency”^90^.

We identified a set of markers for detecting qLSCs that may allow identifying and perhaps purifying qLSCs. What is the functional role of these proteins? Do they control quiescence and/or maintain stemness? What controls their specific expression by qLSCs? Gpha2 (Glycoprotein hormone alpha-2) was found here to be uniquely expressed by qLSCs but its role is still unclear. Transgenic overexpression of Gpha2 in mice showed no gross phenotype^91^. Interestingly, we report a drastic reduction of Gpha2 levels in cases where the niche was disturbed, for example, when LSCs were disconnected from the niche and grown *in vitro*, or following immune cell repression or absence *in vivo*. In both cases, quiescence was lost, suggesting that Gpha2 may be regulated by T cells and controls quiescent cell state. By contrast, the interferon-induced transmembrane protein 3 (Ifitm3) was still expressed in the absence of immune cells *in vivo* and *in vitro*, suggesting that it is regulated by other niche means, not via a T cell-induced pathway, and that it does not control cell proliferation. Interestingly, both Ifitm3 and Cd63 reside in cellular endo membranes^92^. Moreover, Ifitm3 is known for its antiviral specific functions^93^, providing protection against entry and replication of several types of viruses including the COVID-19 virus^71^. Ifitm3 was shown to be highly expressed in embryonic, neural and mesenchymal SCs, suggesting that it may confer anti-viral activity to qLSCs. Interestingly, it was shown that Ifitm3 was involved in B cell related leukemia^94^ and control embryonic germ cells^95^. Such evidence hints that ifitm3 may provide protection and control important functions of SCs. No major developmental disorder was reported regarding ifitm3 knockout mice, although the cornea or LSCs have not been investigated in that study^96^. Emb (Embigin), an adhesion molecule that marked qLSCs (Fig. S3a), was shown to regulate hematopoietic SC quiescence and localization^97^. Furthermore, Emb was linked with quiescence of hair follicle SCs^55^ while Gpha2 and Ifitm3 were shown to be expressed in epidermal SCs in the G0 state^98^. Future studies will be needed to illuminate the role of these genes in SC homeostasis and pathology, and to test whether their manipulation can contribute to advancing LSC therapy.

In summary, this report provides a useful atlas that uncovers the main corneal epithelial cell populations, capturing the signature and the niche of quiescent and activated LSC states. These data open new research avenues for studying the mechanisms of cell proliferation and differentiation as well as the applications of LSCs in regenerative medicine.

## Methods

### RNA-seq protocol and analysis

Ten eyes of 2.5 months old Krt15-GFP mice were enucleated, the limbus (with marginal conjunctiva and peripheral cornea) was dissected (∼0.5 mm), tissues were pooled and incubated in trypsin (X10 Biological Industries) for 10 min 37°C 300 μl. Supernatant was collected into 10 ml (RPMI (Biological Industries) 10% chelated fetal calf serum). Trypsinization was repeated for 10 cycles adding fresh trypsin in each cycle. Cell suspension was centrifuged (8 min at 300g), re-suspended and filtered using Cell strainer (Danyel Biotech) to achieve 1200 cells/μL. An RNA library was produced according to the 10x Genomics protocol (Chromium Single Cell 3’ Library & Gel Bead Kit v2, PN-120237) using 18000 input cells. Single cell separation was performed using the Chromium Single Cell A Chip Kit (PN-120236). The RNAseq data was generated on Illumina NextSeq500, 150 bp paired-end reads, high-output mode (Illumina, FC-404-2005) according to 10X recommendations read 1-26 bp and read 2-98bp. Cell ranger (version 2.2.0) was used for primary analysis. Transcripts were mapped to the mm10 reference genome with the addition of the eGFP gene sequence. Downstream analyses including clustering, classifying single cells and differential expression analyses were performed using R package – Seurat (version 2.3.4)^46^. Low-quality cells (>5% mitochondrial UMI counts, or <200 or >8000 expressed genes^99^) and genes (detected in <3 cells) were excluded, and eventually 14,941 genes across 4,738 cells were analyzed.

In silico analysis of non-epithelial cell markers confirmed the absence (>0.1%) of corneal endothelial cells (Clrn1, Pvrl3, Gpc4, Slc9a7, Slc4a4, Grip1, Hrt1d, Irx2), corneal stromal keratocytes (Vim, Thy1, S1004a4), goblet cells (Muc2, Muc5ac, Muc5b), melanocytes (Dct, Mitf, Ptgs, Tyrp1, Melana, Tyr), T cells (Cd3d, Cd3e, Cd4, G2mb), B cells (Cd19, Cd79a, Cd79b). The only non-epithelial cells identified in our dataset were six cells that expressed MHC II (H2-Ab1^+^), 4 of them were Cd11c^+^/Cd207^+^ or Cd11c^+^/Cd39^+^ dendritic-like cells, 2 others were Cd45^+^/Adgre^+^ or Cd45^+^/Cd64^+^ macrophage-like. Analysis of pan epithelial genes (expression of Krt14, Krt12, Cdh1, Epcam and Cldn7) confirmed that most (>99%) cells were ocular epithelial cells. For pathway analysis, a list of differentially expressed genes between chosen clusters was analyzed using WebGestalt^56^. Over-Representation Enrichment Analysis (ORA) was used for pathway analysis with KEGG functional database.

### Animals, tissue preparation and staining

Animal care was according to the ARVO Statement for the Use of Animals in Ophthalmic and Vision Research. R26R-Brainbow^2.1^ (#013731), K14-Cre^ERT2^ (#005107), Krt15-GFP (#005244) and UBC-CreERT2 (#008085) were from Jackson Laboratories (Bar Harbor, ME). To induce Cre recombinase activity, 4 mg/day Tamoxifen (T5648, Sigma, St. Louis, MO) dissolved in corn oil was intraperitonealy injected (200 μl) for 3-4 consecutive days, as previously reported^29 30^. NOD/SCID, SCID, Nude, and BALB/c were from Envigo RMS Ltd (Israel). Dexamethasone (Sigma) was dissolved in PBS (0.5%) and administered as drops (15 µl) on ocular surface of anesthetized (2% Isoflurane) mice, for 10 minutes every 12 hours for 4 consecutive days. For wounding, mice were anesthetized (2% Isoflurane) and injected intramuscular with analgesics Buprenorphine (0.03 mg/ml, 50 µl). Central corneal wounding (2 mm diameter) was performed using an ophthalmic rotating burr (Algerbrush) under fluorescent binocular. The wounded corneas were stained (1% fluorescein), and the wounded area was imaged and quantified (NIS-Elements analysis D). For CD4^+^FoxP3^+^ regulatory T cell depletion, subconjunctival injection of 30 µl (250 µg) anti-Cd25 antibody (Clone:PC61, antiCd25 IL-2Rα) or rat IgG (control) (BioCell *InVivo*MAb)^100^ was performed.

Single intraperitoneal injection of 200 μl (7.5 mg/ml) EdU (Sigma) was performed and 6 hours later tissues were processed. For pulse-chase, EdU was injected every 12 hours for 14 consecutive days and tissues processed after additional 30-days. Isolated corneas were fixed (2% PFA, 1 hour) and stained (Click-iT, Invitrogen) according to the manufacturer’s instructions followed by whole mount staining protocol (described above) for other markers staining. For the calculation of length cycle (Tc) 1.7 mg/ml IdU (Abcam) was injected and 1.5 hours later, 7.5 mg/ml EdU was injected and tissues harvested 30 minutes later. The calculation of length cycle (Tc) was calculated according to formula described by^62^

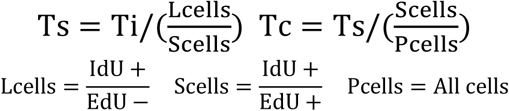

### Staining, imaging and quantifications

For paraffin sections, eyes were fixed (overnight, 4% formaldehyde), washed briefly with phosphate buffered saline (PBS), incubated in increasing ethanol, Xylene and paraffin (60°C) and then embedded in paraffin blocks, as previously described^33^. Sections (5 μm) were prepared, antigen retrival performed with unmasking solution (Vector Laboratories, H3300), blocked (0.2% Tween20 and 0.2% gelatin, 2 hours), incubated with primary antibody (overnight, 4°C), secondary antibodies (Invitrogen, 1:500, 2 hours) followed by 4′,6-diamidino-2-phenylindole (DAPI) staining, and mounting (Thermo Scientific). For H&E (Sigma) staining, sections were rehydrated, incubated with Hematoxylin (7 minutes), washed (tap water, 15 minutes), stained with Eosin (3 minutes), dehydrated, incubated with Xylene (15 minutes) and mounted (Thermo Scientific). For frozen sections, eyes were fixed (4 hours, 4% formaldehyde), washed briefly (PBS), incubated in 30% sucrose (overnight) and embedded in optimal cutting temperature compound (OCT). Sections (5 μm) were fixed (4% paraformaldehyde), permeabilized (0.1%, TritonX-100, 20 minutes), blocked (1% BSA, 1% gelatin,2.5% Normal goat serum, 2.5% Normal donkey serum, 0.3% TritonX-100, 1 hour), incubated with primary antibody (overnight, 4°C), secondary antibodies (1:500, 1 hour), DAPI and mounting (Thermo Scientific). For whole mount, corneas were isolated, fixed (2% formaldehyde, 2 hours, room temperature), permeabilized (0.5% Triton, 5 hours), blocked (0.1% TritonX-100, 2% Normal donkey serum, 2.5% BSA, 1 hour), primary antibody (overnight, 4°C on a shaker), secondary antibodies (1:500, 1 hour), followed by DAPI, tissue flattening under a dissecting binocular and mounting (Thermo Scientific), as previously described^33^. ISH was performed according to the manufacturer’s instructions for frozen tissues (LGC Biosearch Technologies). All probe sets contained ≥ 28 oligonucleotides each (20-mer). Hybridization mixture contained 125 nM of each oligonucleotide probe mix (Krt4 (059668, set 48), Gpha2 (024784, set 28), Atf3 (026628, set 48), Krt12 (020912, set 47)).

Staining conditions were adjusted for each antibody used for immunofluerescent staining on paraffin (IF-P) or frozen section (IF-Fr) or wholemount cornea (IF-Wmt) or cultured cells (IF-C) or Western blot (WB). We used the following conditions: Krt4 (Abcam ab183329, IF-Fr/IF-P/IF-Wmt 1:100), Krt8 (Abcam ab53280, IF-Fr/IF-P/IF-Wmt 1:10), Krt6a (BioLegend 905702, IF-P 1:100), Gpha2 (Santa cruz sc-390194, IF-Wmt 1:1000, IF-P 1:100, IF-Wmt more recommended), Cd63 (Santa cruz sc-5275, IF-P/IF-Wmt 1:100), Ifitm3 (Abcam ab15592, IF-P/IF-Wmt/ 1:10,000, IF-Fr/IF-C/WB 1:1000), Atf3 (Santa cruz sc-188, IF-P 1:100), Mt1-2 (Abcam ab12228, IF-Wmt 1:100 *permabilization for 2 hours), Krt12 (Abcam ab185627, IF-Fr/IF-P/IF-C/WB 1:1000), Krt15 (Santa Cruz sc-47697, IF-Fr/IF-P/IF-C/WB 1:1000), Krt17 (Cell signaling 4543S, IF-P 1:100), Krt19 (Abcam ab5265, IF-P 1:100), Krt14 (Millipore CBL197, IF-P 1:1000), P63 (Santa Cruz sc-8431, IF-C 1:100), Cd31 (Abcam ab231436, IF-Wmt 1:100), Lyve-1 (Abcam ab14917, IF-Wmt 1:100), Cd3e (BioLegend 100311, IF-Wmt 1:100 APC conujugated), Cd8 (BioLegend 100706, IF-Wmt 1:100), Cd45 (BioLegend 103102, IF-Wmt 1:100), Cd19 (BD Pharmingen 553786, IF-Wmt 1:100), GFP (Abcam ab13970, IF-Fr 1:1000), IdU (Abcam ab187742, IF-Wmt 1:100), Ki67 (Abcam ab16667, IF-Fr/IF-P/Wmt 1:100), Gapdh (Cell signaling #2118, 1:1000 WB), Krt13(Abcam ab16112, IF-P 1:500)

Images were acquired using 20X/0.8 M27 Plan-Apochromat objective on LSM 880 airyscan laser scanning confocal system (Zeiss, Oberkochen, Germany) or by Nikon Eclipse NI-E upright Microscope using Plan Apo λ 20x objective. For computerized quantification analyses we used Nis-elements Analysis D software, the outer limbus was marked by Gpha2 or Cd63 staining, the adjacent 100 µm (or entire inner limbus if K15-GFP used) of the inner limbus was calculated, while 100 µm peripheral cornea was quantified and 5 different field were calculated from each cornea (n=3-5).

### *In vitro* experiments

Human limbal rings from cadaveric corneas were obtained under the approval of the local ethical committee and declaration of Helsinki. Cells cultivation, differentiation, transfection, Western blot and real time polymerase chain reaction (qPCR) analyses were performed as previously described^101^. We used esiRNA against EGFP (EHUEGFP, Sigma), IFITM3 (EHU222801, Sigma) or GPHA2 (EHU036941, Sigma) and cells were collected 5-6 days after transfection. Primers for qPCR were: GAPDH (GCCAAGGTCATCCATGACAAC, CTCCACCACCCTGTTGCTGTA), IFITM3 (CTGGGCTTCATAGCATTCGCCT, AGATGTTCAGGCACTTGGCGGT), KRT15 (GACGGAGATCACAGACCTGAG, CTCCAGCCGTGTCTTTATGTC), KRT12 (TGAATGGTGAGGTGGTCTCA, TTTCAGAAGGGCAAAAAGGA).

### Clone size and number analysis

As described previously^27 23 60^, one of the main predictions of the neutral drift dynamics model is that the average number of cells in a given clone as a function of time is given by, 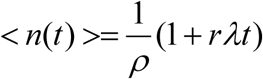 where *ρ* is the fraction of basal cells that divide, 2r is the probability for symmetric division and *λ* is the division rate. The number of clones is expected to decrease with time as, 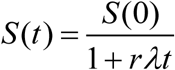. Thus, the relation between the clone size and clone number is given by 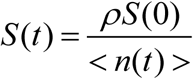 Liner fit of the clone size was performed using Matlab linear regression model. Estimating the clone size cumulative distribution was done using empirical CDF estimation using Matlab.

### Statistical analysis

Data are presented as means +/- SD. T-test, ANOVA (analysis of variance) followed by Bonferroni test, or Kolmogorov-Smirnov test were performed using GraphPad Prism software, as indicated in legends, to calculate p-values. Differences were considered to be statistically significant from a p-value below 0.05.

## Acknowledgement

We thank L. Linde, S. Galili and their team for the scRNA-seq experiment, D. Aberdam and Y. Bar Onn for critical reading. RSF received funding from European Union’s Horizon 2020 research & innovation programme (828931), the ISRAEL SCIENCE FOUNDATION (1308/19 and 2830/20), Rappaport Family Institute for Research in Medical Sciences, NIH-exploratory R21 (800040).

## Author Contribution

A.A. and A.A.-L. were involved in conceptual and experimental design, performed and interpreted the experiments, prepared the figures, analyzed data and participated in the manuscript writing; N.T., S.D., L.S., S.B. performed experiments, prepared the figures and participated in the manuscript writing; S.H.-P., W.N, J.I., G.B.-D. performed experiments and provided data; B.T., E.B. and N.K. provided materials and participated in discussions and manuscript writings; Y.S. and R.S.-F. was involved in conceptual and experimental design, data interpretation, and manuscript writing. All authors approved the manuscript.

## Figure Legends

**Figure S1: Exploration and validation of the identity of cell populations**. Corneas of adult 2-3 months old Krt15-GFP mice were used for analysis. (A) Whole-mount immunostaining of Krt8 (upper panel) and bright field image of the same zone (lower panel). (B) Hierarchical clustering of gene expression data from scRNA-seq. (C) Violin plots showing the expression of the indicated keratin genes each cluster. (D) Heatmap presentation showing the average expression of the indicated cell cycle genes by each cluster. Immunostaining of wholemount cornea (E-F, H) or cornea sections (G) using antibodies against the indicated markers. The dashed squares shown in (E) are shown (DAPI only) at higher magnification in (F). (H) EdU pulse/chase experiment described in Fig. 3e-f that was performed to detect slow cycling cells. The square zone on the right shows enlarged view of the dashed region on the left. Arrow shows EdU^+^/Krt17^+^ limbal epithelial cells at the outer limbus. Scale bars are 50 μm. Abbreviations: Conj., Conjunctiva; Peri., Periphery.

**Figure S2: Prediction of pathways enriched in clusters 3 or 4**. In silico analysis performed to predict pathways that were enriched in cluster 3 or 4 (in comparison to all other clusters) using the Webgestalt algorithm.

**Figure S3: Differential expression of selected families of genes**. Heatmap presentation shows the average expression of the indicated genes across clusters. Genes enriched in cluster 3 or cluster 4 are shown in A and B, respectively, and selected family-related genes are shown in (C).

**Figure S4: Comparative analysis of proliferation and dynamic limbal epithelial cell populations**. (A) Genes that are differentially expressed between clusters 3 and 4 were identified by WebGestalt/KEGG enrichment analysis. (B-C) Schematic representation of R26R-Confetti; UBC-CreERT2 double transgenic mice (B) and the potential outcome of tamoxifen induction of Cre recombinase (C). (D-E) Lineage tracing was induced by tamoxifen injection in 2-month old UBC-Cre; Brainbow^2.1^ mice that were sacrificed at the indicated times post induction, corneas were Immunostaining for Gpha2 and typical confocal pictures are shown (the same pictures without Gpha2 channel are shown in Fig. 1A-B). (F) The number of clones in the same area as a function of time for different regions. (G) One minus the cumulative distribution function of the clone size for different time points in different regions (compare to Fig 2H which shows the scaled distributions). Scale bars, 50 μm. Abbreviations: Conj., Conjunctiva; Peri., Periphery.

**Figure S5: Role of T cells as qLSC niche cells**. Whole-mount immunostaining of immunodeficient mice (SCID (A-B), NOD SCID (C), and Nude (E)) and their controls (BALB/c). Tissues were stained for the indicated markers and confocal images were purchased. Z-stack confocal pictures of the basal epithelial layer (epithelium) or anterior stroma (stroma) is shown in (A). (D) H&E staining of the corneal epithelium of immuneodefficient (SCID) mice and their control showing mild epithelial thickening. (F) The ocular surface of adult mice was topically treated with dexamethasone (or vehicle as control) for 4 days. Confocal images of corneas stained with the indicated antibodies are shown. Scale bars are 50 μm. Abbreviations: Peri., Periphery.

