## Supplemental for "Capturing limbal epithelial stem cell population dynamics, signature, and their niche"

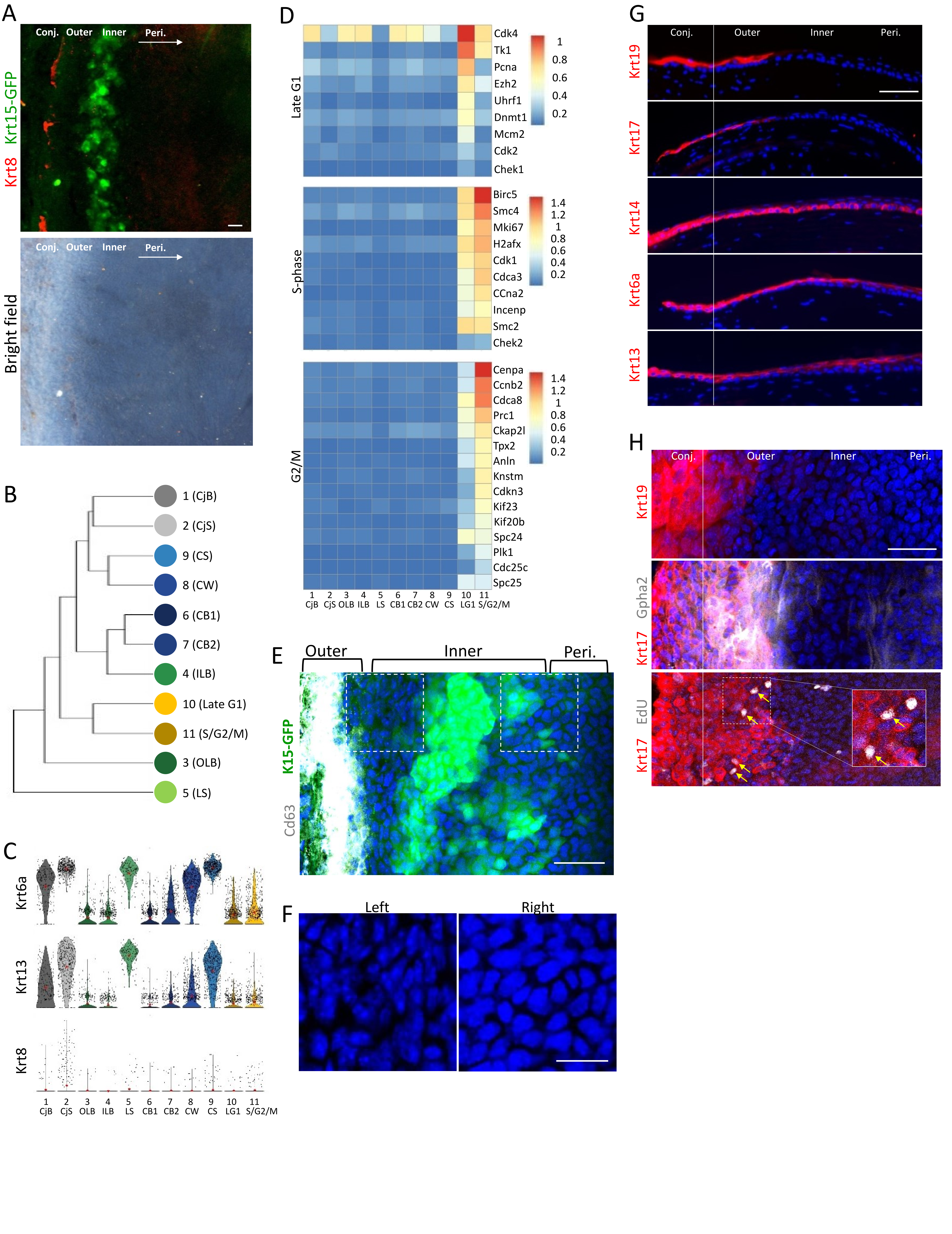


Fig S1: Exploration and validation of the identity of cell populations

A

B

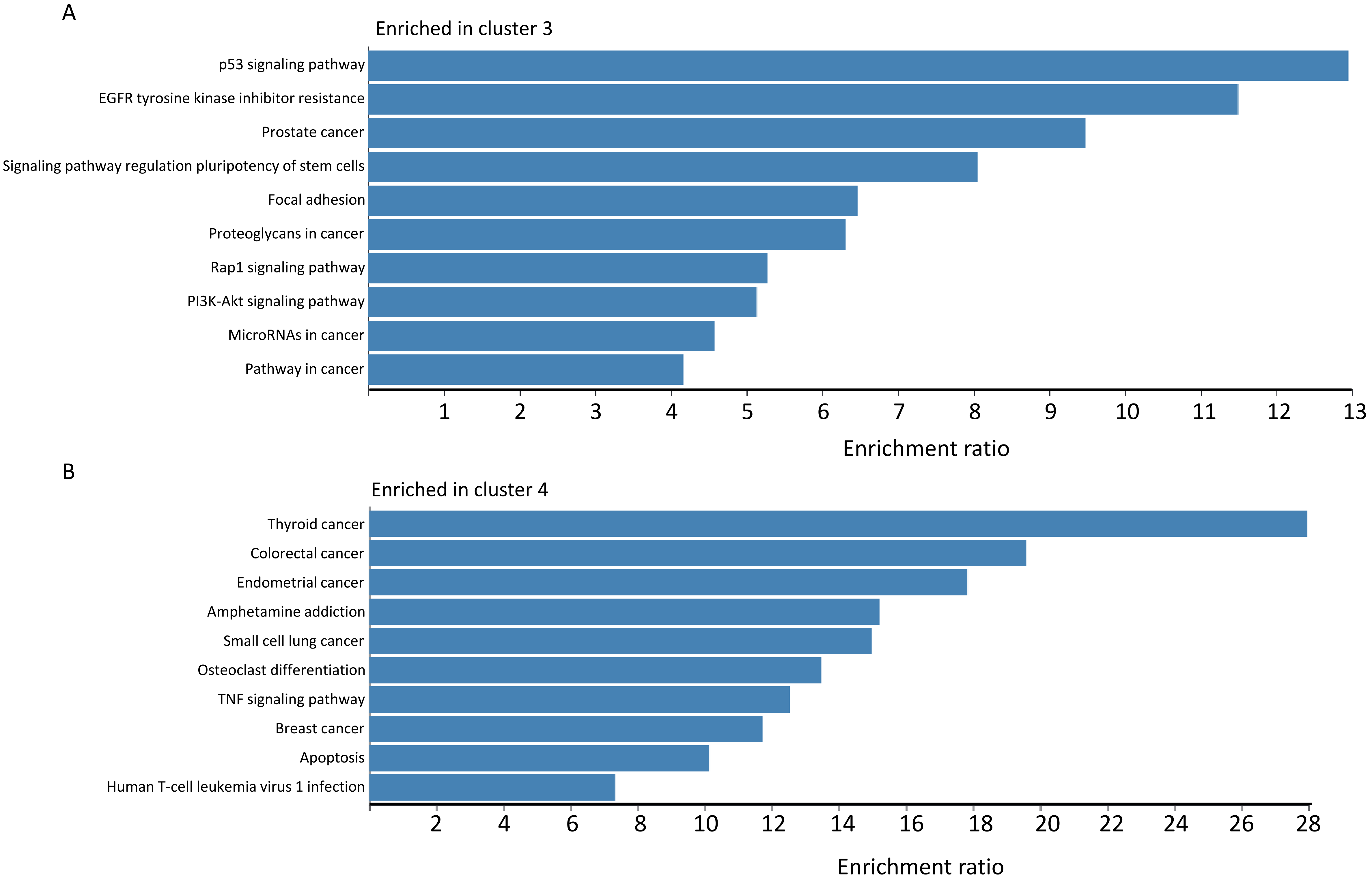


Fig S2: Prediction of pathways enriched in clusters 3 or 4

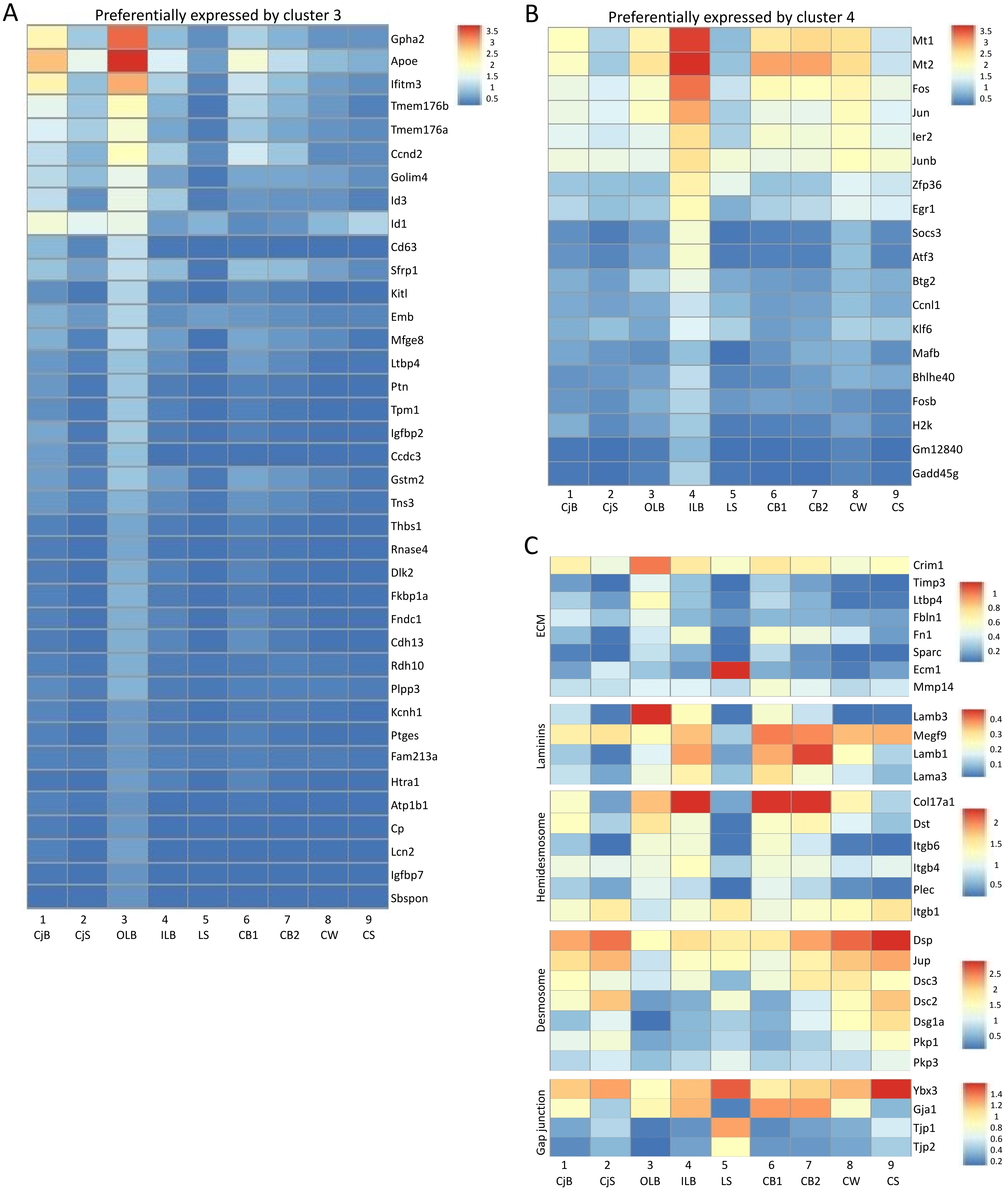


Fig S3: Differential expression of selected families of genes

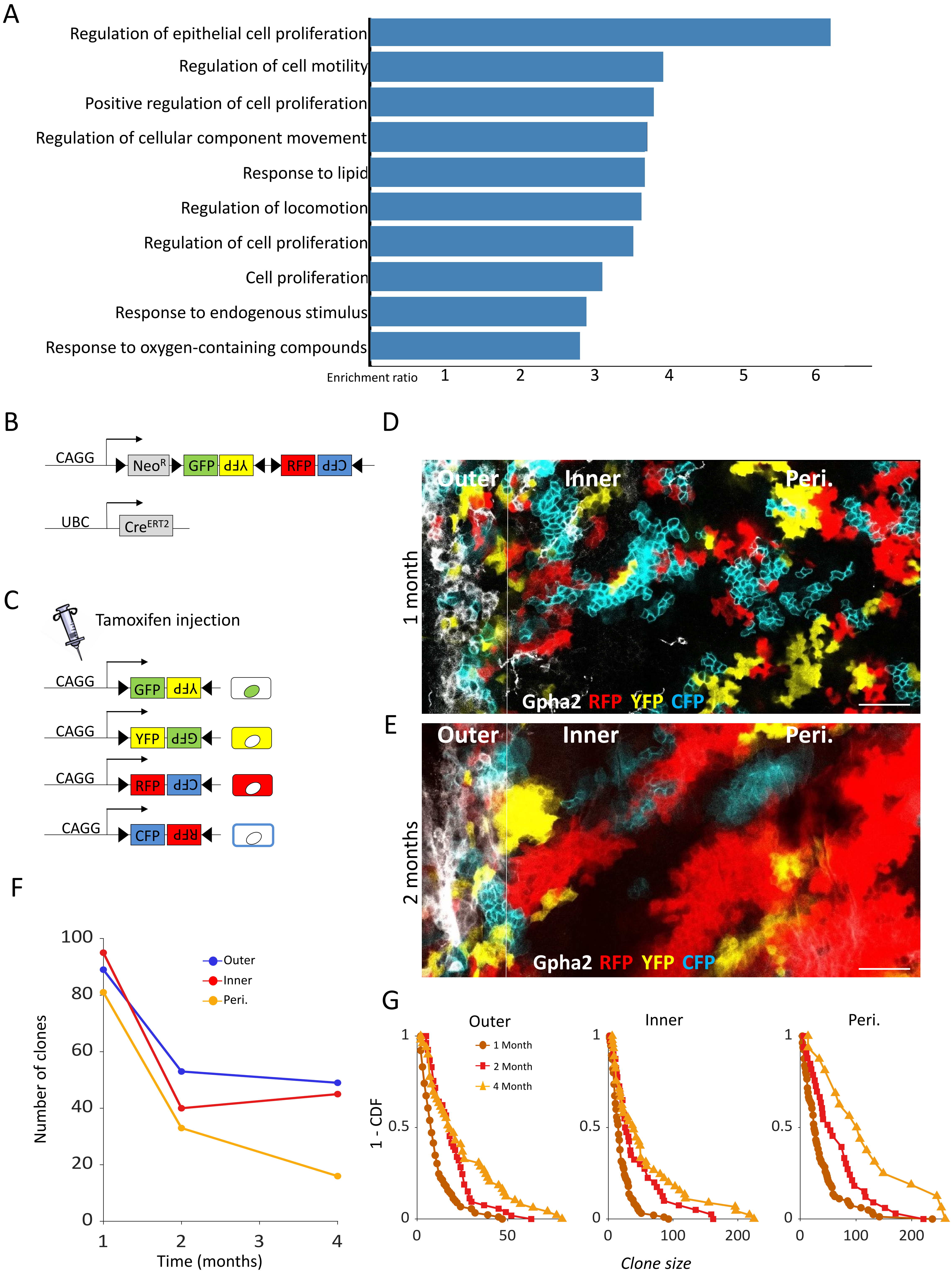


Fig S4: Comparative analysis of proliferation and dynamic limbal epithelial cell populations

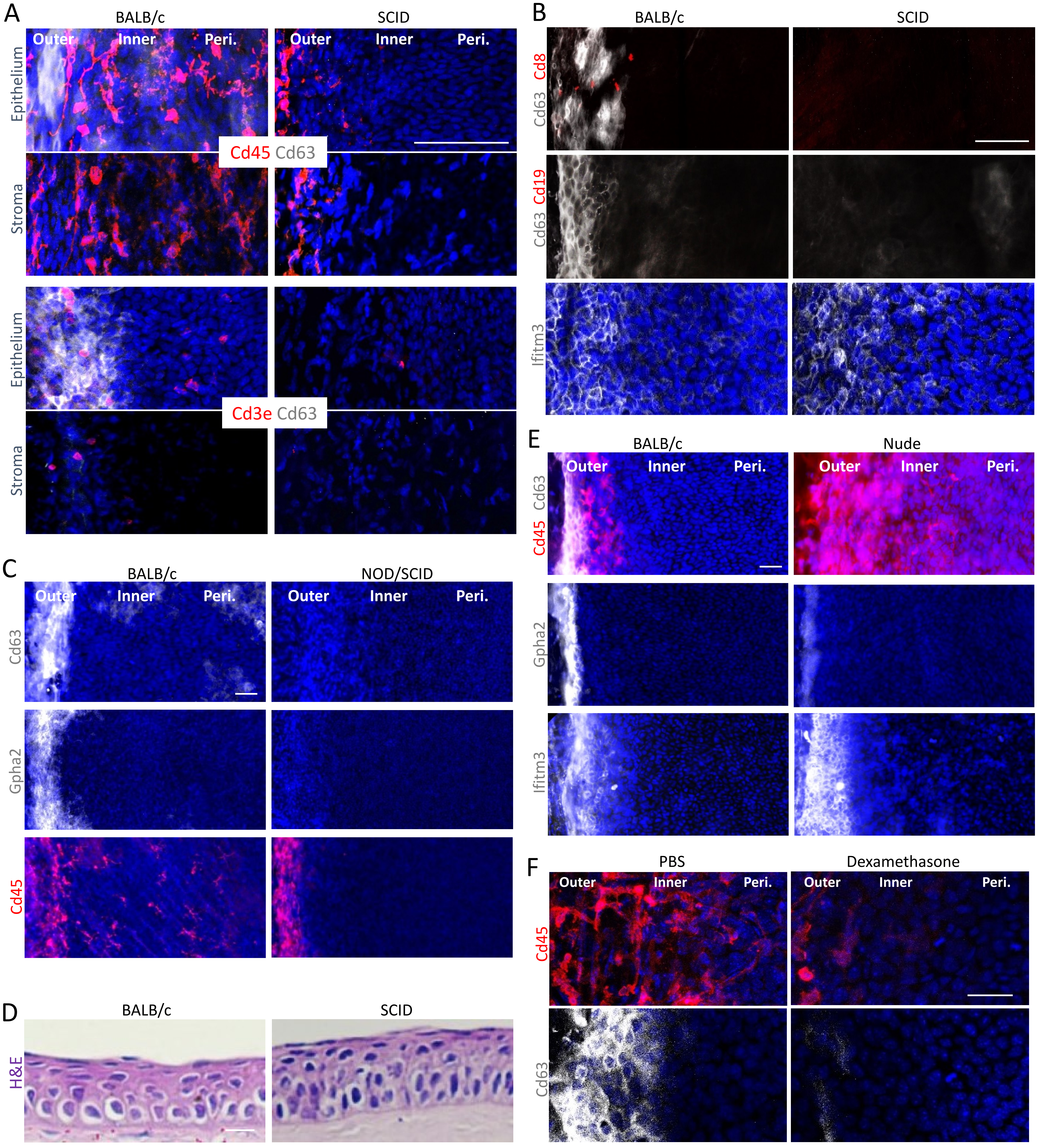


Fig S5: Role of T cells as qLSC niche cells
